# α-catenin mechanosensitivity as a route to cytokinesis failure through sequestration of LZTS2

**DOI:** 10.1101/2023.08.25.554884

**Authors:** Yuou Wang, Alex Yemelyanov, Christopher D. Go, Sun Kim, Jeanne M. Quinn, Annette S. Flozak, Phuong M. Le, Shannon Liang, Anne Claude-Gingras, Mitsu Ikura, Noboru Ishiyama, Cara J. Gottardi

## Abstract

Epithelial cells can become polyploid upon tissue injury, but mechanosensitive cues that trigger this state are poorly understood. Using α-catenin (α-cat) knock-out Madin Darby Canine Kidney (MDCK) cells reconstituted with wild-type and mutant forms of α-cat as a model system, we find that an established α-cat actin-binding domain unfolding mutant designed to reduce force-sensitive binding to F-actin (α-cat-H0-FABD^+^) can promote cytokinesis failure, particularly along epithelial wound-fronts. Enhanced α-cat coupling to cortical actin is neither sufficient nor mitotic cell-autonomous for cytokinesis failure, but critically requires the mechanosensitive Middle-domain (M1-M2-M3) and neighboring cells. Disease relevant α-cat M-domain missense mutations known to cause a form of retinal pattern dystrophy (α-cat E307K or L436P) are associated with elevated binucleation rates via cytokinesis failure. Similar binucleation rates are seen in cells expressing an α-cat salt-bridge destabilizing mutant (R551A) designed to promote M2-M3 domain unfurling at lower force thresholds. Since binucleation is strongly enhanced by removal of the M1 as opposed to M2-M3 domains, cytokinetic fidelity is most sensitive to α-cat M2-M3 domain opening. To identify α-cat conformation-dependent proximity partners that contribute to cytokinesis, we used a biotin-ligase approach to distinguished proximity partners that show enhanced recruitment upon α-cat M-domain unfurling (R551A). We identified Leucine Zipper Tumor Suppressor 2 (LZTS2), an abscission factor previously implicated in cytokinesis. We confirm that LZTS2 enriches at the midbody, but discover it also localizes to tight and tricellular junctions. LZTS2 knock-down promotes binucleation in both MDCK and Retinal Pigmented Epithelial (RPE) cells. α-cat mutants with persistent M2-M3 domain opening showed elevated junctional enrichment of LZTS2 from the cytosol compared α-cat wild-type cells. These data implicate LZTS2 as a mechanosensitive effector of α-cat that is critical for cytokinetic fidelity. This model rationalizes how persistent mechano-activation of α-cat may drive tension-induced polyploidization of epithelia post-injury and suggests an underlying mechanism for how pathogenic α-cat mutations drive macular dystrophy.

## INTRODUCTION

Cytokinesis failure is rapidly emerging as a means to new cell and tissue physiologies rather than an unproductive glitch (Bailey et al., 2021; Fox et al., 2020; Peterson & Fox, 2021; Schoenfelder & Fox, 2015). Cardiomyocytes become binucleated through a developmentally timed defect in cytokinetic furrow progression (Engel et al., 2006; Normand & King, 2010; Soonpaa et al., 1996). This leads to terminally differentiated cells that are larger than their mononuclear (diploid) counterparts (Patterson et al., 2017), presumably to support the long term function of these continuously contracting actin-myosin sarcomere units (Karbassi et al., 2020; Orr-Weaver, 2015; Øvrebø & Edgar, 2018). Macrophages can also become multinuclear upon exposure to certain bacteria by eliciting Toll-Like Receptor signals that antagonize cytokinesis. This leads to formation of giant polyploid macrophage “granulomas” that are transcriptionally distinct for managing chronic infections (Herrtwich et al., 2016). Even simple epithelia can become polyploid after tissue injury due to failed cytokinesis (Cao et al., 2017; Lazzeri et al., 2019), where increased DNA copies may enable adaptation to an injured state that facilitates repair (Schoenfelder & Fox, 2015). Unique to epithelial barriers, injury-induced epithelia polyploidization and consequent hypertrophy is advantageous, as larger cells manifest less junctional surface area per unit of epithelial area covered, leading to decreased permeability in an otherwise leaky-barrier environment (Cohen et al., 2018; Losick et al., 2013). Thus, cytokinesis failure appears to be a “feature, not a bug” of the mitosis paradigm, leading to a diversity of beneficial outcomes.

If benefits of cytokinesis failure are clear, the way cells fail to successfully complete this process are just emerging. Excessive integrin-extracellular matrix adhesive tension along the wound front appears to be one cause of cytokinesis failure, where basal stress fibers can interfere with contractility of the actin-myosin cytokinetic ring leading to a cell with two nuclei (Uroz et al., 2018, 2019). Curiously, the adherens junction (AJ) component, alpha-catenin (α-cat), is a tension-sensitive actin-binding protein that sustains missense mutations linked to macular dystrophies (Saksens et al., 2016). Remarkably, a forward genetic screen in mice aimed at modelling the earliest stages of disease suggests pathogenesis initiates through progressive multinucleation of retinal pigment epithelial cells. How α-cat dysfunction leads to multinucleation in this system, however, remains unknown.

α-cat mechanosensitivity and AJ function depend on the conformations and binding activity of α-cat’s N-terminal, Middle and C-terminal regions (Ishiyama et al., 2013; Rangarajan & Izard, 2012). The N-terminal domain comprises two 4-helical bundles, where the former binds ý-catenin and associates with cadherin adhesion receptors (Pokutta & Weis, 2000). The C-terminal 5-helical bundle shows allosteric binding to F-actin, where force-dependent alteration of α-helical (H) regions H0 and H1 favors high affinity binding of H2-5 to actin filaments, leading to “catch-bond” behavior (Buckley et al., 2014; Ishiyama et al., 2018; Wang et al., 2022; Xu et al., 2020). The middle (M)-region of α-cat (comprising three 4-helical bundles (M1-3)) can undergo sequential, force-dependent unfurling events to recruit α-cat M-domain binding partners (Barrick et al., 2018; Kim et al., 2015; Li et al., 2015; Maki et al., 2018; Pang et al., 2019; Seddiki et al., 2018; Terekhova et al., 2019; Thomas et al., 2013; Yao et al., 2014; Yonemura et al., 2010). But the features of adherens junction (AJ) organization and dynamics impacted by distinct α-cat unfolding states, and the mechanosensitive α-cat binding partners linked to AJ organizational states are only just emerging (Cho et al., 2022; Donker et al., 2022a; Ishiyama et al., 2018a; Matsuzawa et al., 2018; Monster et al., 2021a; Noordstra et al., 2023; Sakakibara et al., 2020; Sarpal et al., 2019; Sheppard et al., 2023; Twiss et al., 2012; van den Goor & Miller, 2022). In the current study, we show that alterations in α-cat binding to F-actin or its mechanosensitive middle (M)-domain are sufficient to drive cytokinesis failure and sustain binucleated cells within an epithelial monolayer. We further credential Leucine Zipper Tumor Suppressor 2 (LZTS2), an understudied scaffold protein previously implicated abscission (Sudo & Maru, 2007, 2008), as a novel conformation-sensitive proximity partner of α-cat, where persistent recruitment to AJs interferes with LZTS2 ring formation at the midbody. These findings demonstrate that LZTS2 is a mechanosensitive effector of α-cat critical for abscission fidelity in epithelia and suggest that tension-dependent changes on α-cat and AJ may contribute to the phenomenon of epithelial cell polyploidization post-injury for means of effective barrier repair.

## RESULTS

### Persistent coupling of α-cat to cortical actin enhances epithelial cell binucleation

To understand how α-cat binding to actin is regulated by force, Ishiyama et al engineered an α-cat F-actin-binding domain (FABD) unfolding mutant (RAIM è GSGS mutation in α-helix 0) with the goal of attenuating force-dependent α-cat binding to actin, where this mutant leads to enhanced F-actin binding and epithelial monolayer strength compared with wild-type α-cat (Ishiyama et al., 2018). By reconstituting this mutant, α-cat-H0-FABD^+^, into α-cat knock-out Madin Darby Canine Kidney (MDCK) cells, we generalized this phenomenon to another epithelial cell type, showing that partial loss of the α-cat catch bond mechanism leads to stronger epithelial sheet integrity with greater co-localization between the α-cat-H0-FABD^+^ mutant and actin. This mutant, however, also interferes with more dynamic processes such as epithelial wound-closure (Ishiyama et al., 2018)(Wood et al., 2023). Since AJ of α-cat-H0-FABD^+^ -restored MDCK cells do not show constitutive accessibility to the α18 monoclonal antibody M2-domain epitope compared with WT α-cat, we reason this mutant enhances static but interferes with dynamic cell-cell adhesive behaviors through persistent engagement with lower tension cortical actin networks (Wood et al., 2023).

Having established α-cat-H0-FABD^+^ as a tool to interrogate contributions of α-cat catch-bond activity to dynamic adhesive processes (i.e., a “force-desensitized” actin-binding form of α-cat), we discovered that the α-cat-H0-FABD^+^ mutant -restored MDCK α-cat KO cells were associated with significantly higher rates of binucleation compared with WT-α-cat-restored cells (Fig. 1A-D). This effect was enhanced along epithelial wound-fronts (Fig. 1E), a process associated with increased tension on the cadherin-catenin complex (Borghi et al., 2012; Donker et al., 2022; Monster et al., 2021). Based on these data, we hypothesized that altered α-cat mechanosensitivity might causally contribute to the generation and/or maintenance of polyploid cells that emerge along wound fronts and are critical for epithelial barrier repair (Cao et al., 2017).

**Fig.1:**
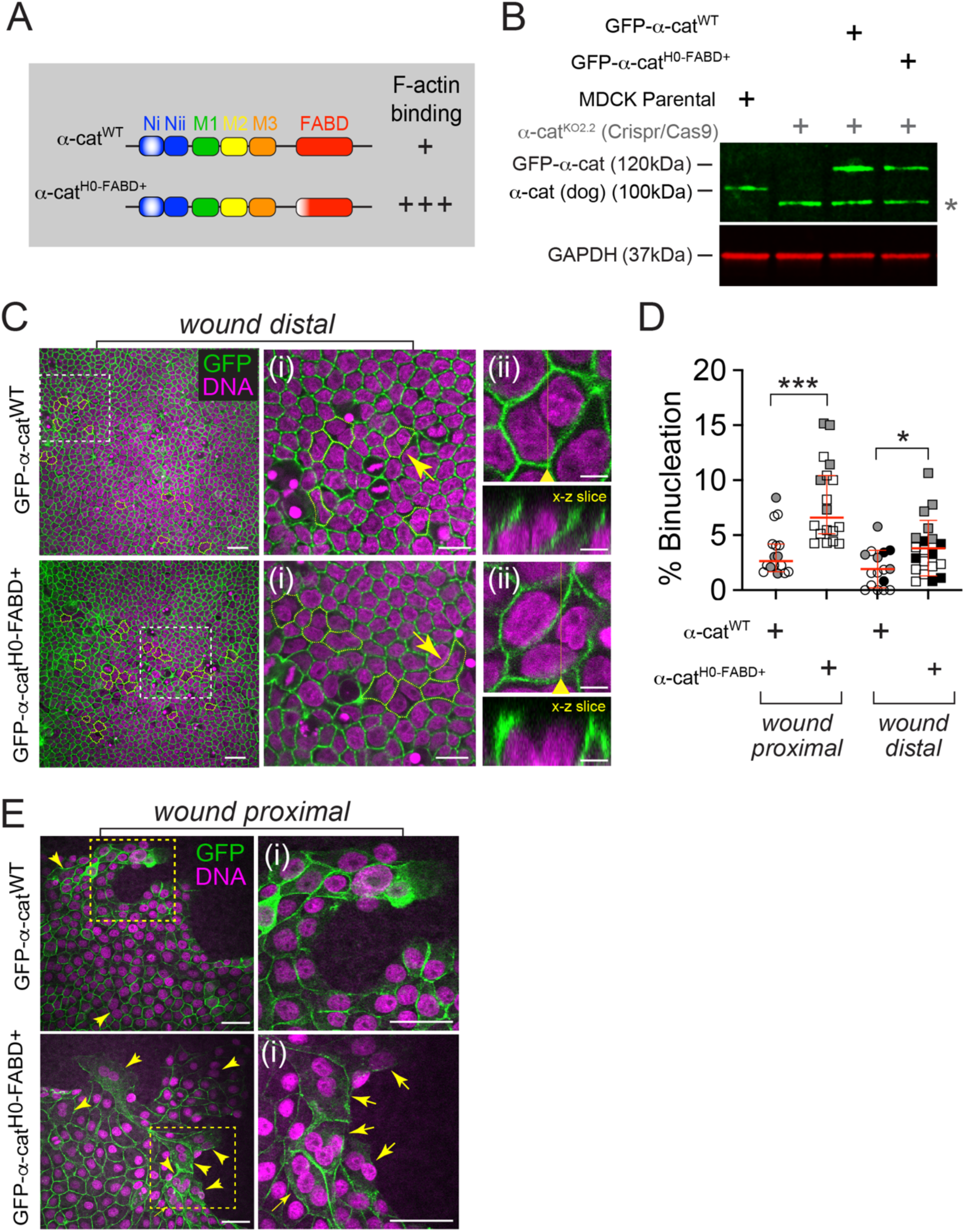
Persistent coupling of α-cat to F-actin enhances binucleation rate. **(A)** Schematic of α-cat domains based on crystal structure. α-cat^H0-FABD+^ replaces 4 amino acids (RAIM>GSGS) in the first kinked helix of α-cat’s five helical bundle actin-binding domain (whitening of red domain), compromising normal force-gated binding of a-cat to F-actin. This generates an α-cat that binds F-actin more strongly (+++) than α-cat (+) via in vitro pelleting assays (Ishiyama, 2018). **(B)** Immunoblot of MDCK KO2.2 cells transfected with GFP-tagged versions of WT α-cat or an F-actin-binding-domain (FABD) unfolding mutant (a-cat^H0-FABD+^). GAPDH was used as loading control. Asterisk (*) marks an α-cat fragment generated by our CRISPR-targeting strategy via an alternative translational start site (L C Bullions et al., 1997). As this α-cat isoform lacks the ý-cat binding region and fails to robustly localize to cell-cell contacts (Wood et al., 2023), it does not likely contribute to the cytokinesis interference pathway in this study. **(C)** Representative immunofluorescence images of MDCK α-cat KO cells restored with GFP-α-cat^WT^ (Top) or GFP-α-cat^H0-FABD+^ (Bottom). GFP-α-cat (green), nuclei (magenta), binucleated cells (outlined, yellow dashed line). Scale bar, 30μm. Rectangular regions framed in white-dashed boxes are enlarged to show detail (right, i). Scale bar, 15μm. Yellow arrows denote binucleated cells chosen for orthogonal views (right, ii). Yellow arrowhead shows plane of x-z slice. Scale bar, 5μm. **(D)** Graph shows percentage of wound distal versus wound proximal cells displaying binucleation. p<0.05 (*) and <0.001 (***) by unpaired t test. Symbols denote measurements from distinct fields of view; different colors reflect biological replicates (n=3). **(E)** Leading edge of MDCK α-cat KO cells expressing GFP-α-cat^WT^ (Top) or GFP-α-cat^H0-FABD+^ (Bottom) 40-hr post scratch wound. GFP-α-cat (green), nuclei (magenta). Yellow arrows denote binucleated cells. Scale bar, 40μm. The rectangular regions framed in yellow are enlarged to show detail (right, i). Scale bar, 40μm.

### Persistent coupling of α-cat to actin promotes binucleation via cytokinesis failure

The increased association between the α-cat-H0-FABD^+^ mutant and MDCK cell binucleation could occur via two distinct routes: cytokinesis failure or cell-cell fusion, two common routes of epithelial cell polyploidization after injury (Bailey et al., 2021; Dehn et al., 2021; Gjelsvik et al., 2019; Nandakumar et al., 2020; Øvrebø & Edgar, 2018; Peterson & Fox, 2021). To investigate the route to polyploidization observed above, we performed live imaging of GFP-α-cat-WT or GFP-α-cat-H0-FABD^+^ cells co-expressing the histone 2B-red fluorescent protein reporter (H2B-RFP) (Fig. 2; Movie 1, left). Interestingly, mitosis took much longer in α-cat-H0-FABD^+^ (46.00 min) than WT α-cat-restored MDCK cells (32.17 min). Telophase and cytokinesis phases were substantially extended in α-cat-H0-FABD^+^ (14.71 min) compared with α-cat-WT cells (4.61 min), and relative to other mitotic phases (Fig. 2A, B). Remarkably, α-cat-H0-FABD^+^ cells also showed significantly higher rates of cytokinesis failure, particularly during the latest stage of cytokinesis, abscission (Fig. 2A, asterisk, 2C-graph; Movie 1, right). In all cases where binucleated cells formed during imaging, an intervening mitotic event was captured, suggesting cytokinesis failure rather than cell-cell fusion is the major route to binucleation in α-cat-H0-FABD^+^ -expressing cells. Thus, enhanced coupling of α-cat to cortical actin can interfere with cytokinetic fidelity.

**Fig. 2:**
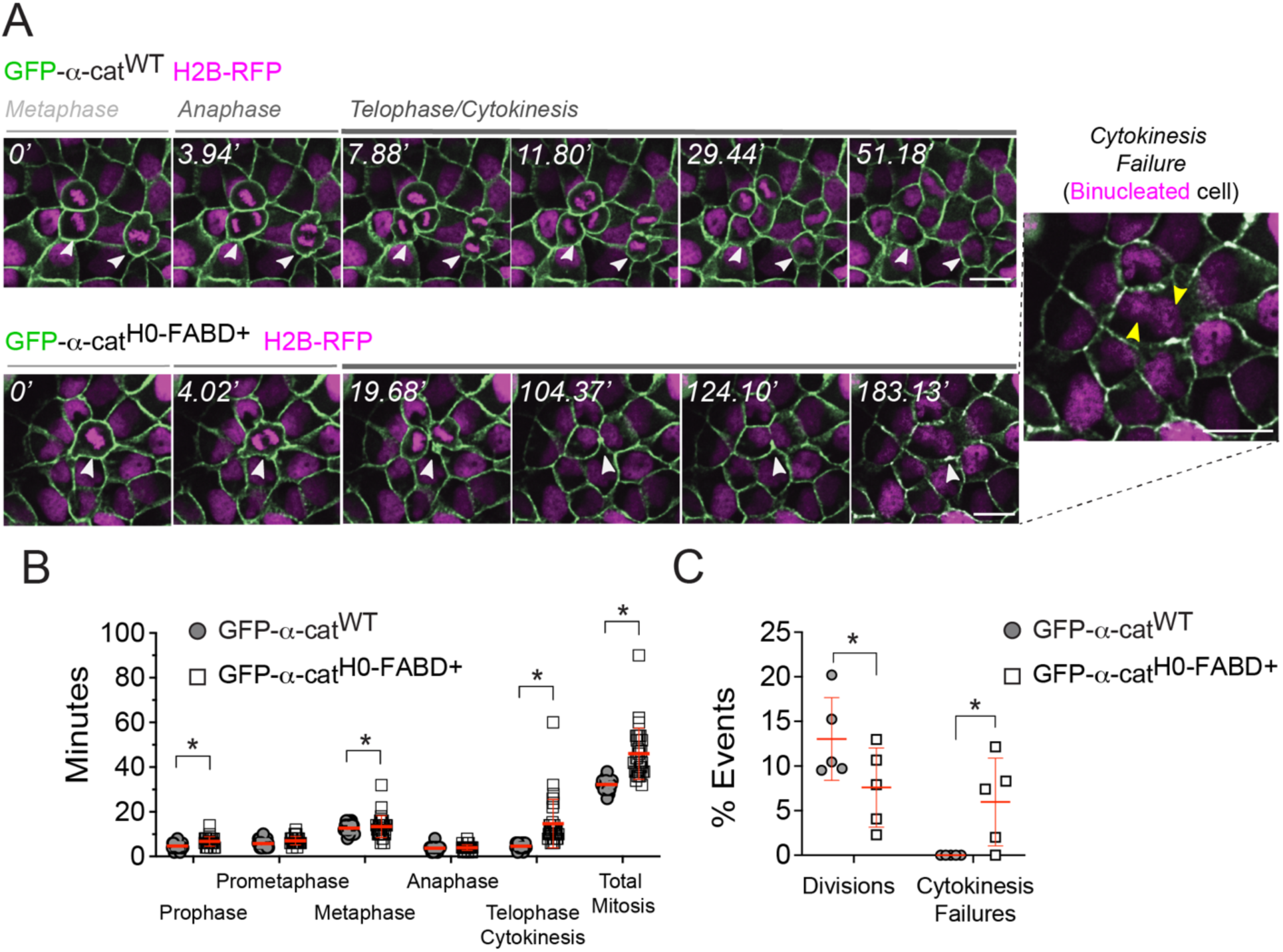
Persistent coupling of α-cat to F-actin promotes binucleation via cytokinesis failure. **(A)** Live imaging analysis of MDCK α-cat KO cells restored with α-cat^WT^(Top) or α-cat^H0-FABD+^(Bottom) during mitosis phases (metaphase, anaphase, telophase and cytokinesis;). Time stamp in minutes (‘) shown in white; GFP-α-cat (green), nuclei (magenta), white arrowheads indicate dividing cells. A binucleated cell after cytokinesis failure is shown (inset with yellow arrowheads shows two nuclei in one cell). See also Movie 1. Scale bar, 30μm **(B)** Quantification of α-cat^WT^ or αCat^H0-FABD+^ cells (N = 30); *p < 0.05 by unpaired t-test. **(C)** Graph shows percentage of cell divisions and cytokinesis failures over 5 fields of view; *p<0.05, unpaired t-test.

### α-cat-H0-FABD^+^-induced cytokinesis failure does not occur in single cells

Cell division can be regulated both autonomously and non-autonomously (Fededa & Gerlich, 2012; Herszterg et al., 2013; Monster et al., 2021b; Pinheiro et al., 2017). To understand if α-cat regulates cytokinesis through neighboring cells, we characterized mitosis at the single cell level. Interestingly, while α-cat-H0-FABD^+^ -restored MDCK single cells showed significant mitotic-phase lengthening, telophase and cytokinesis preceded normally with no evidence of abscission failure (Fig. S1 A, B). Even α-cat KO parental cells that remained attached through the mitotic-rounding/re-adhesion sequence showed similar kinetics of mitosis with no cytokinesis failures, although the number of events captured was much fewer (see Discussion for caveat). Similar results were observed in the E-cadherin/catenin-deficient epidermoid cell line, A431D, where enforced α-cat-H0-FABD^+^ expression failed to impact any stage of mitosis compared with WT-α-cat (Fig. S1C, D). Together, these data suggested α-cat can only interfere with cytokinetic fidelity from within the cadherin-catenin complex, and in a manner that depends on neighboring cell-cell contacts.

### α-cat-H0-FABD^+^-induced cytokinesis failure requires the mechanosensitive M-domain

Evidence that WT α-cat and α-cat-H0-FABD^+^ show similar M2-domain accessibility to the α18 monoclonal antibody raises the possibility that enhanced coupling to cortical actin can occur in the absence of α-cat M2-domain unfurling (Wood et al., 2023). To distinguish whether α-cat-H0-FABD^+^ interferes with cytokinesis through binding to F-actin alone, or via α-cat M-domain conformation-sensitive functions consequent to ABD binding to F-actin, we engineered WT α-cat and α-cat-H0-FABD^+^ mutants lacking the entire M-domain (Fig. 3A). Interestingly, both α-cat-ΔM and α-cat-ΔM H0-FABD^+^ progressed through mitosis similarly to WT-α-cat (Fig. 3B-C). However, telophase lengthening and cytokinesis failures associated with the α-cat-H0-FABD^+^ mutant were completely avoided when the M-domain was removed from this mutant (Fig. 3D, α-cat-ΔM H0-FABD^+^). These data suggest persistent coupling of α-cat to cortical actin alone is insufficient to interfere with cytokinesis, and that the conformationally sensitive M-domain is required for interference.

**Fig. 3.**
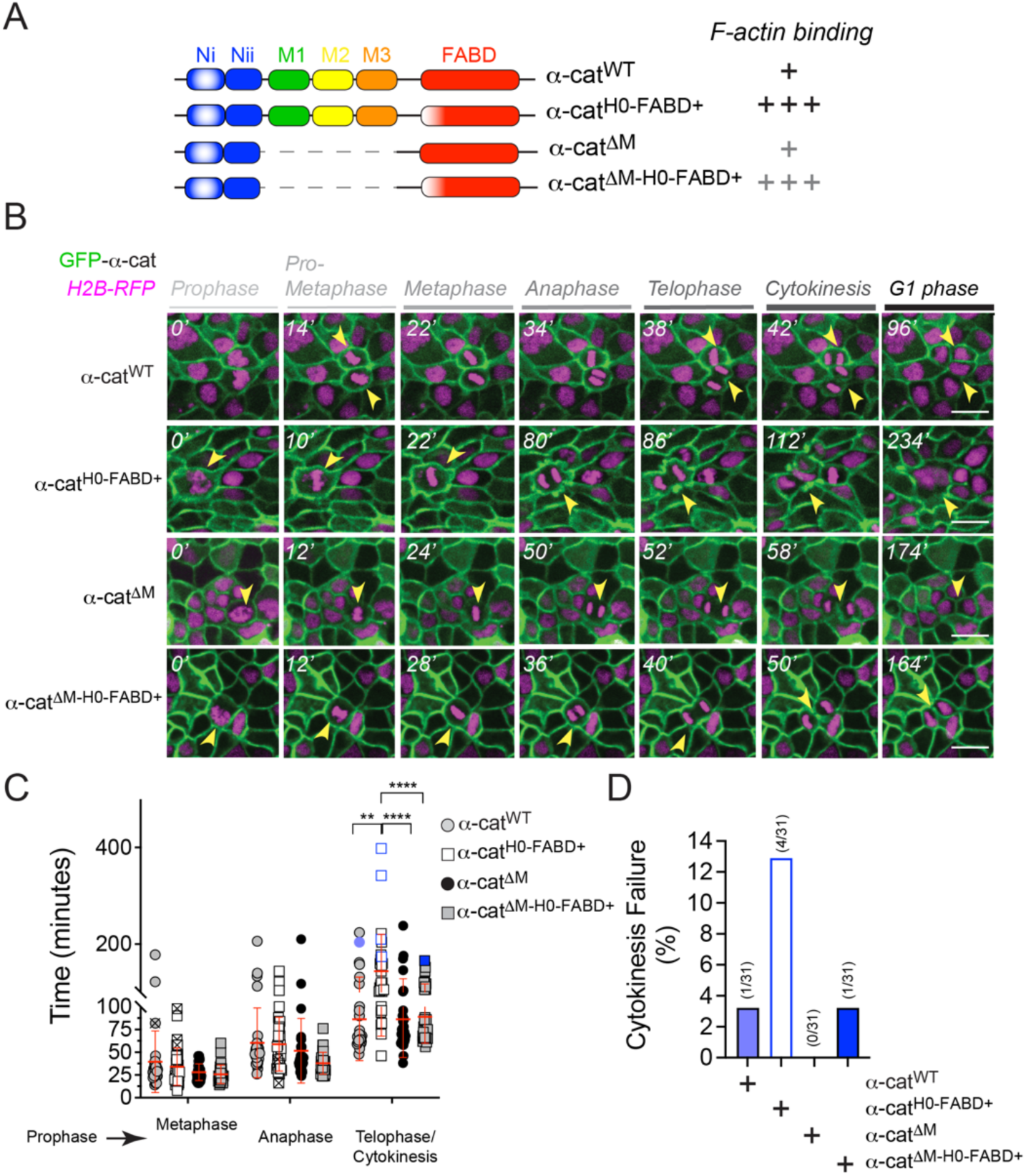
α-cat^H0-FABD+^ -induced cytokinesis failure requires the mechanosensitive M-domain. **(A)** Schematic of α-cat mutants used for analysis. (**B**) Live imaging analysis of MDCK α-cat KO cells restored with α-cat^WT^, α-cat^H0-FABD+^, α-cat^ýM^ and α-cat^ýM-H0-FABD+^ during mitosis phases (gray). Time stamp in minutes (‘, white); yellow arrowheads show dividing cells; GFP-α-cat (green), nuclei (magenta). Scale bar, 50μm **(C)** Graph quantification of mitosis phase lengths as minutes from prophase. ** and **** p< 0.01 and 0.001 by ANOVA. **(D)** Graph quantification of cytokinesis failure events.

### α-cat M-domain missense mutants causing eye dystrophy support binucleation

While full length α-cat protein is essential for organismal development (Ishiyama et al., 2018; Sheikh et al., 2006; Torres et al., 1997), M-domain localized missense mutations in the human gene *CTNNA1* are associated with butterfly-shaped pigment dystrophy (BPD), a rare patterned dystrophy of the retinal pigment epithelium that leads to a progressive, age-dependent loss of vision (Saksens et al., 2016; e.g., α-cat-E307K). Remarkably, a forward genetic screen in mice aimed at modelling the earliest stages of this patterned dystrophy identified a mouse *Ctnna1* missense mutant also implicating α-cat M-domain dysfunction in this disease (α-cat-L436P).

Curiously, disease pathogenesis initiates through progressive multinucleation of retinal pigment epithelial cells, raising the possibility that α-cat M-domain dysfunction might causally contribute to this process. To learn whether these M-domain missense mutations could enhance the rate of binucleation in a heterologous epithelial cell system, we stably expressed α-cat-E307K or -L436P mutants in MDCK α-cat KO cells (Fig. 4A,B). While the binucleation rate was only modestly increased in dense filter-grown monolayers, this rate was significantly elevated after scratch-wounding (Fig. 4C,D). Live imaging mature monolayers revealed all binucleated cells were generated after mitotic rounding and furrow regression, consistent with cytokinesis failure (Fig. S2A-C). Although binucleated cells do migrate faster than neighboring mononuclear cells as previously described (Kozyrska et al., 2022), this leader cell behavior was not significantly different between WT or mutant α-cat expressing cells (Fig. S2D-F). These data demonstrate that the increased binucleation rate observed along α-cat missense mutant wound fronts relative to WT is due to differences in cytokinesis failure, rather than similarly enhanced rates of wound-induced cytokinesis failure and faster migration of α-cat-E307K or -L436P mutants.

**Fig. 4:**
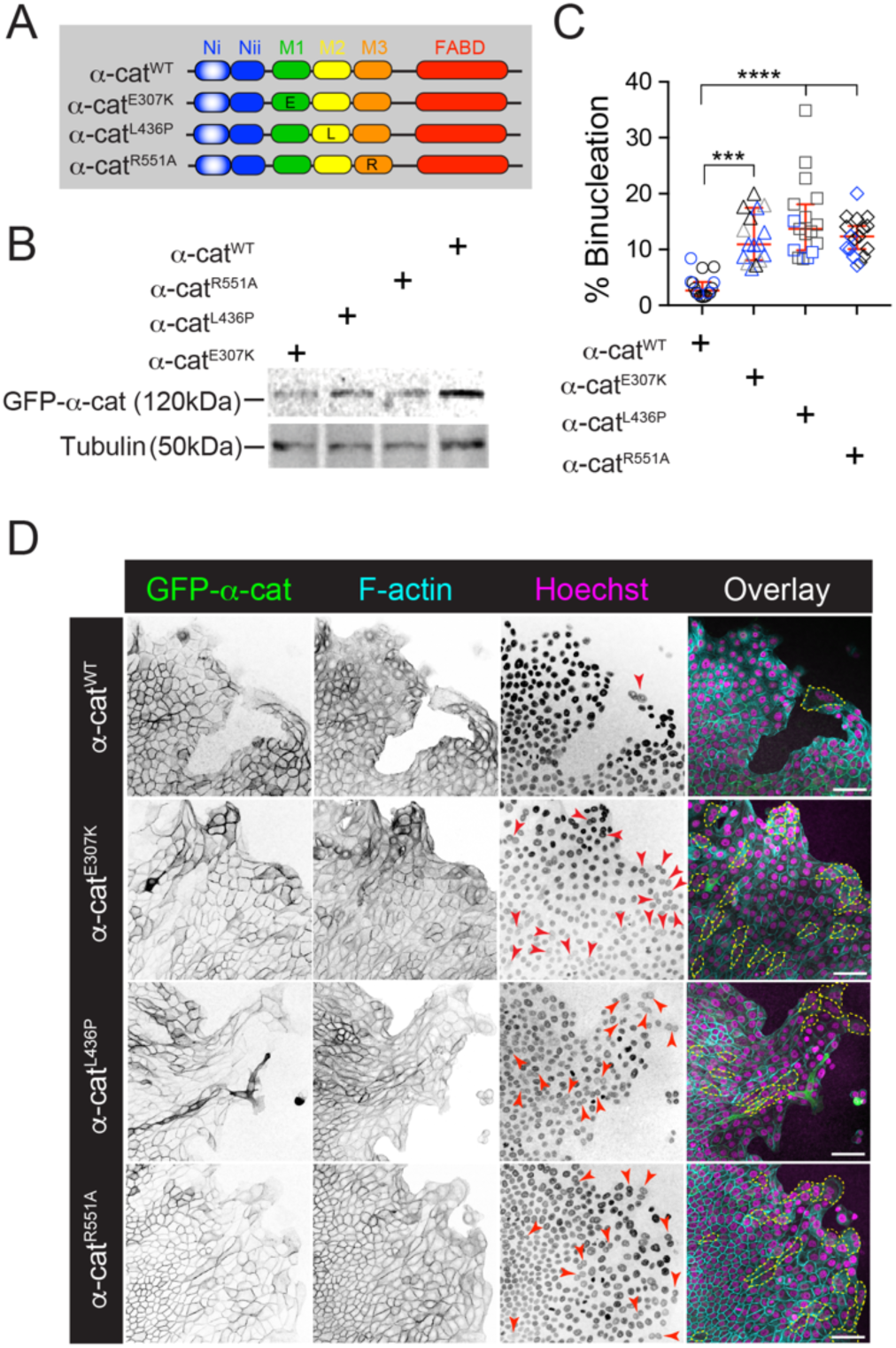
α-cat M-domain missense mutants that cause eye dystrophy promote wound-proximal binucleation. **(A)** Schematic of α-cat missense mutants used for analysis. **(B)** Immunoblots of total cell extracts showing expression levels of α-cat missense mutants. **(C)** Graph quantification of wound proximal binucleation rate (representative images in D). Mean ± SD, two-tailed unpaired t test, ***p<0.001, ****p<0.0001. Data from three independent replicates. **(D)** Inverted-grayscale images showing MDCK α-cat KO cells expressing GFP-α-cat^E307K^, GFP-αCat^L436P^, GFP-αCat^R551A^ are prone to binucleation post-scratch wounding. GFP-α-cat (green), F-actin (Phalloidin; cyan), nuclei (magenta), binucleated cells (outlined, yellow dashed line). Red arrows denote touching nuclei typical to polyploid cells. Scale bar, 50μm.

Lastly, similar cytokinesis failure rates were observed with a previously characterized α-cat M3-domain salt-bridge mutation R551A, which displays a more open M-domain conformation via FRET-based assay (Barrick et al., 2018) (Fig. S2C). While it is currently difficult to predict how E307K or L436P mutations alter α-cat M-domain conformation and mechanosensitivity, evidence these mutants share similar cytokinesis failure phenotypes suggests a common mechanism.

### Identification of α-cat proximity partners sensitive to M-domain unfurling

To identify α-cat conformation-dependent proximity partners potentially relevant to cytokinesis, we inducibly expressed biotin ligase (BirA*)-α-cat chimeric constructs in HEK293 Flp-In T-REx cells using wild-type and R551A salt-bridge disrupting mutant forms of α-cat. After induction of protein expression and biotinylation, we performed streptavidin purification, and analyzed the samples by mass spectrometry (Fig. 5A). High confidence proximal partners with increased binding to the M-domain unfurled R551A mutant over wild-type α-cat were identified using the SAINTexpress algorithm (Fig. 5B). We confirmed vinculin and afadin as α-cat M-domain conformation sensitive binding partners, but selected an understudied protein for further analysis, Leucine Zipper Tumor Suppressor 2 (LZTS2), because of its localization to the midbody and implication in abscission (Sudo & Maru, 2007a, 2008; Thyssen et al., 2006). Below, we credential LZTS2 as a tension-sensitive α-cat binding partner and its requirement for cytokinesis fidelity in MDCK.

**Fig. 5:**
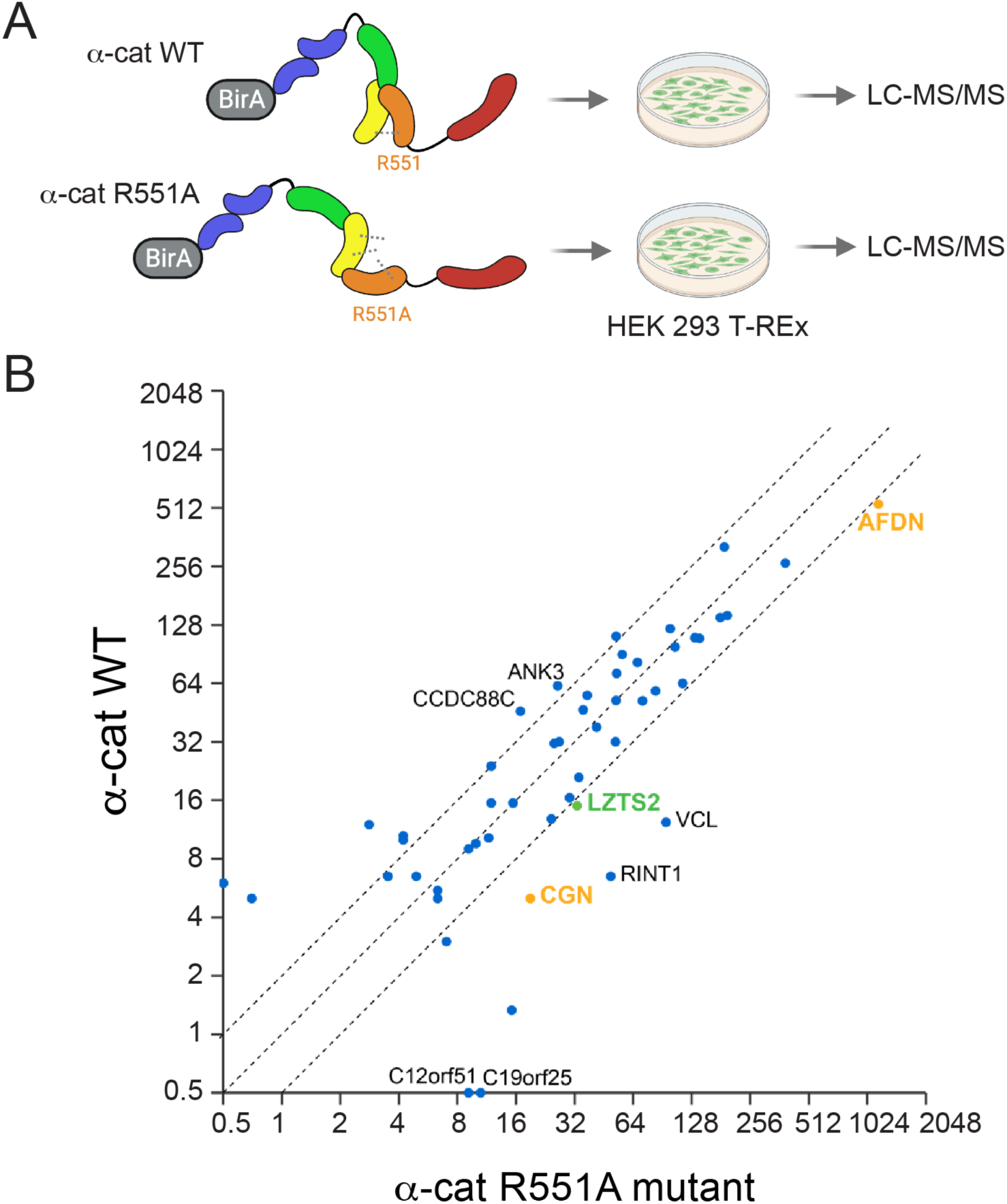
Identification of α-cat proximity partners sensitive to M-domain opening via salt-bridge disruption. **A.** Schematic of experiment: Biotin ligase (BirA*)-tagged α-cat chimeric constructs were expressed in HEK293 Flp-In T-REx cells. Streptavidin purified proteins were analyzed by mass spectrometry. **B**. Scatterplot of α-cat proximity partners with 2-fold greater or reduced enrichment with α-cat salt-bridge disrupted R551A mutant (x-axis) relative to WT α-cat (y-axis). Dotted lines intercepting axes at 1 mark 2-fold enrichment or reduction threshold after normalization.

### LZTS2 localizes to apical and midbody junctions and is required for normal cell division

Recent studies in HEK293 cells reveal that LZTS2 localizes at cell-cell contacts (Go et al., 2021; Baskaran et al., 2021) and the centrosome (Cho et al., 2022). In fully mature filter grown MDCK cells, LZTS2 localizes to apical junctions and the base of primary cilia (Fig. 6A). MDCK cells plated on glass coverslips also show enrichment at multicellular junctions and the midbody (Fig. 6B). To assess roles for LZTS2 in polarized epithelia, we knocked-down LZTS2 using pooled siRNA or shRNA approaches (Fig. 6D). LZTS2 KD MDCKs showed significantly higher binucleation rates compared to negative controls (Fig. 6C&E), consistent with a previous study (Sudo et al., 2007). Similar results were seen in retinal pigmented epithelial (RPE) cells, where higher-efficiency LZTS2 knockdown was more evident by immunoblot analysis, and also led to increased binucleation (Fig. S3A-C). These data show that LZTS2 localizes to cell-cell junctions and midbody structures and its depletion leads to MDCK binucleation.

**Fig. 6:**
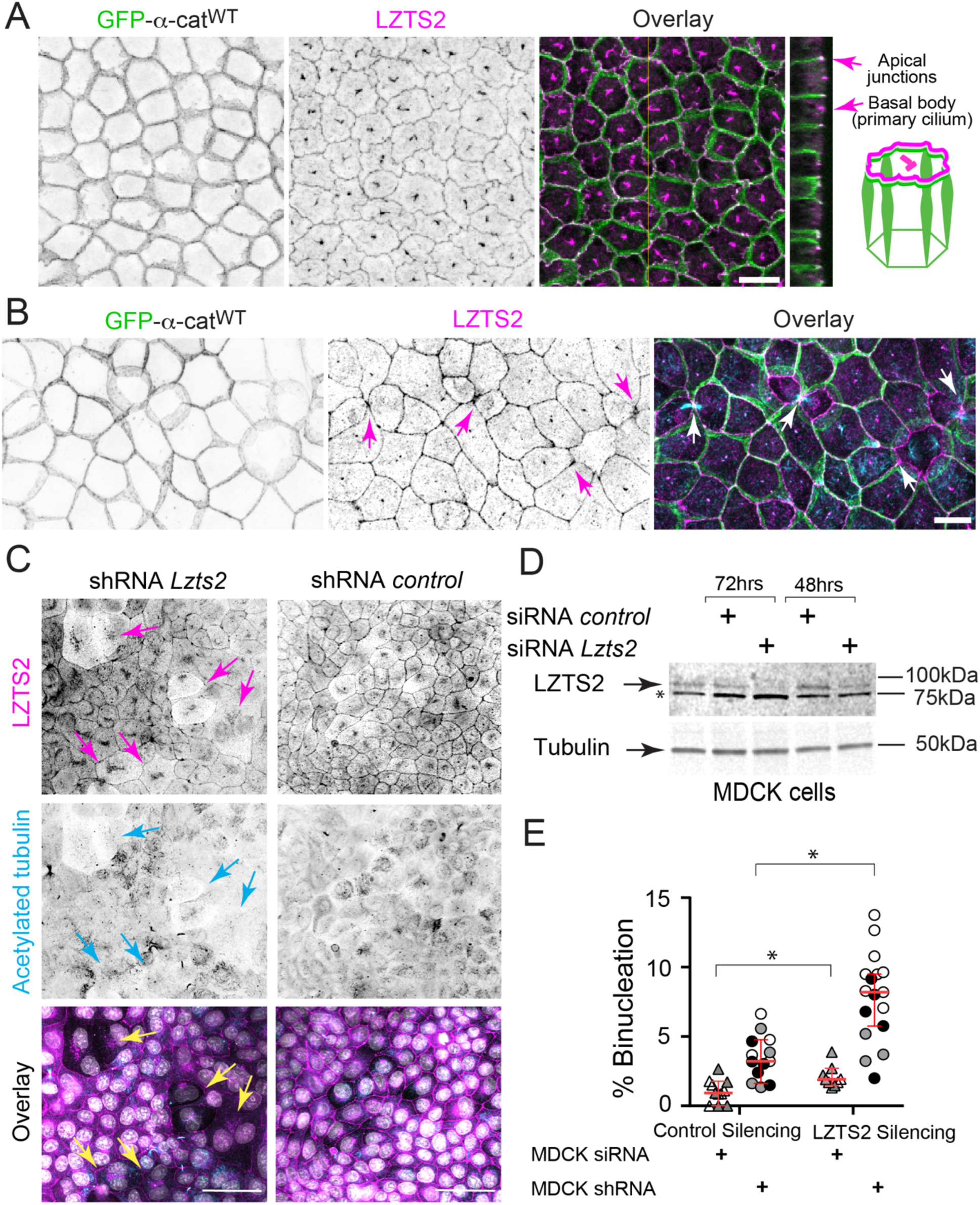
α-cat proximity partner, LZTS2, localizes to apical junctions, base of primary cilia and midbody and is required for successful cell division in MDCK cells. **(A)** Confocal *en face* image (maximum z-projection; inverted grayscale) and orthogonal view images of MDCK cells on Transwell filters showing LZTS2 localizes to apical junctions and basal body (primary cilium), with schematic illustration. α-cat (green) and LZTS2 (magenta). Scale bar, en face 10μm; orthogonal scale, 3μm. **(B)** Confocal images of MDCK cells on coverslips showing LZTS2 is enriched at midbodies during cell abscission (magenta arrows) and tricellular junctions in mitosis (magenta arrowhead). α-cat (green), LZTS2 (magenta), acetylated tubulin (cyan). Scale bar, 10μm. **(C)** Confocal images showing transient knockdown of LZTS2 expression using shRNA in MDCK cells promotes binucleation (colored arrows). LZTS2 (magenta), Acetylated tubulin (cyan). Scale bar, 50μm. Representative images from two independent experiments (n=2) are shown **(D)** Immunoblots of total cell extracts showing partial knockdown of LZTS2 expression using siRNA (48hrs or 72hrs after transfection). Asterisk (*) denotes non-specific band refractory to knock-down and possibly contributing to Golgi localization. **(E)** Percentage of multinucleated MDCK cells upon transient knockdown of LZTS2 expression using either shRNA or siRNA. Two-tailed unpaired t test, *p<0.05. ***p<0.001. Graph indicates mean ± SD. Data from two biological experiments.

### α-cat-1′M1 preferentially recruits LZTS2 to cell contacts over wild-type and other α-cat mutants

Since LZTS2 recruitment to α-cat is M-domain conformation sensitive (Fig. 5) and LZTS2 depletion leads to elevated binucleation (Fig. 6), we reasoned the higher binucleation rate associated with α-cat M-domain and FABD+ mutants might be due to enhanced sequestration of LZTS2 away from the midbody, leading to abscission failure. To address this, we first looked for evidence of bulk LZTS2 sequestration to cell-cell junctions with depletion from the cytoplasmic pool, however, neither α-cat M-domain R551A salt-bridge-disrupting mutant nor missense mutants associated with eye dystrophy showed obvious differences in LZTS2 junction recruitment (Fig. S4). Remarkably, a set of broader α-cat mutants lacking either M1, M2,3 or Nii domains proved more revealing, where the former showed clear evidence of LZTS2 sequestration to cell-cell junctions with a depleted cytosolic pool relative WT α-cat or other mutants (Fig. 7). Since M1 and M2-M3 domains are held together in a closed conformation by salt-bridge interactions (Barrick et al., 2018), deletion of M1 is reasoned to expose M2-M3 helices independently of tension. Thus, M2-M3 domain unfurling appears to “trap” LZTS2 at cell-cell contacts. This effect is consistent with, and quantitatively stronger than recruitment by the M3-domain R551A single point mutant. These data suggest that persistent mechano-activation of α-cat with M2-M3 exposure can alter the junction versus cytoplasmic distribution of LZTS2 in MDCK cells.

**Fig. 7:**
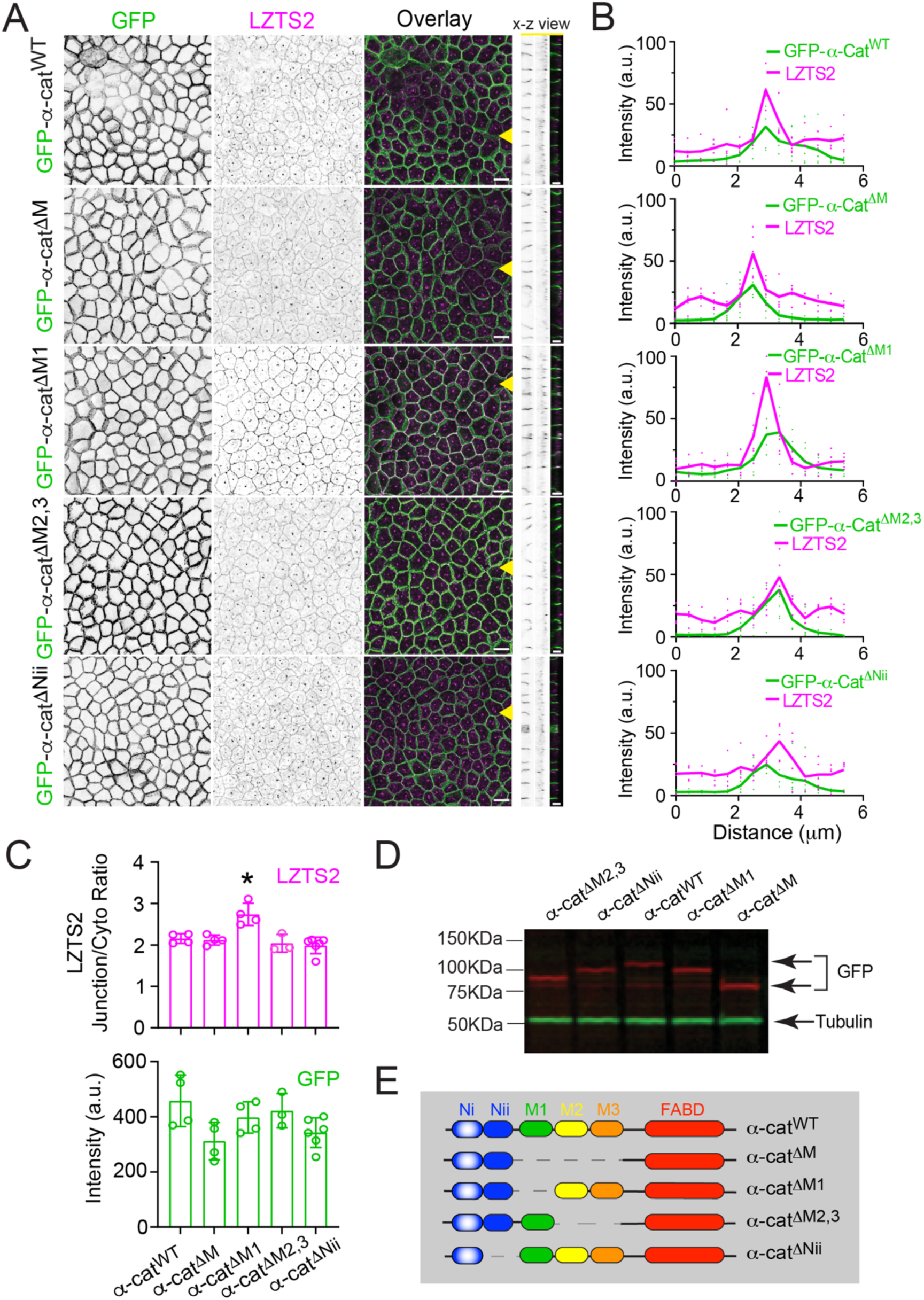
α-cat-1′M1 preferentially recruits LZTS2 to cell contacts over wild-type and other α-cat mutants. **A)** Confocal *en face* views (maximum z-projection; inverted grayscale) and x-z stacks showing α-cat-1′M1 enriches LZTS2 at cellular junctions, α-cat (green), LZTS2 (magenta), Scale bar, x-y view; 10μm, x-z view, 5μm. Representative images from four independent experiments are shown. **(B)** Normalized intensity profiles of GFP-α-cat (green) and LZTS2 (magenta) based on line scans (width: 3 pixels) performed across cytosol to bicellular junctions. Line scan average of 5 junctions/construct. **(C)** Automated quantification of LZTS2 junction/cytosol ratio (upper graph), one way ANOVA, *p<0.05, paired with automated quantification of GFP-α-cat junction intensity (lower graph). Automated quantification based on five randomly selected 90 μm X 90 μm FOVs. **(D)** Immunoblots of total cell extracts showing the expression levels of different GFP-α-cat constructs (red bands) with tubulin loading control (green). **(E)** Schematics of GFP-α-cat constructs.

### LZTS2 recruitment to WT α-cat junctions is actomyosin dependent

To determine if LZTS2 recruitment to cell-cell contacts is actomyosin dependent, we assessed localization under conditions of myosin inhibition. When WT-α-cat-restored MDCK were grown on coverslips, blebbistatin treatment diminished LZTS2 signal, especially along bicellular junctions (Fig. 8A,B). Interestingly, LZTS2 recruitment to α-cat ΔM1-restored MDCK cells was unaffected by blebbistatin, as expected for an α-cat mutant that constitutively exposes M2-M3 helices independently of tension (Fig. 8C,D). These data suggest that junctional recruitment of LZTS2 is tension sensitive, and that the α-cat ΔM1 mutant leads to a persistently enhanced recruitment of LZTS2 to cell-cell contacts.

**Fig. 8:**
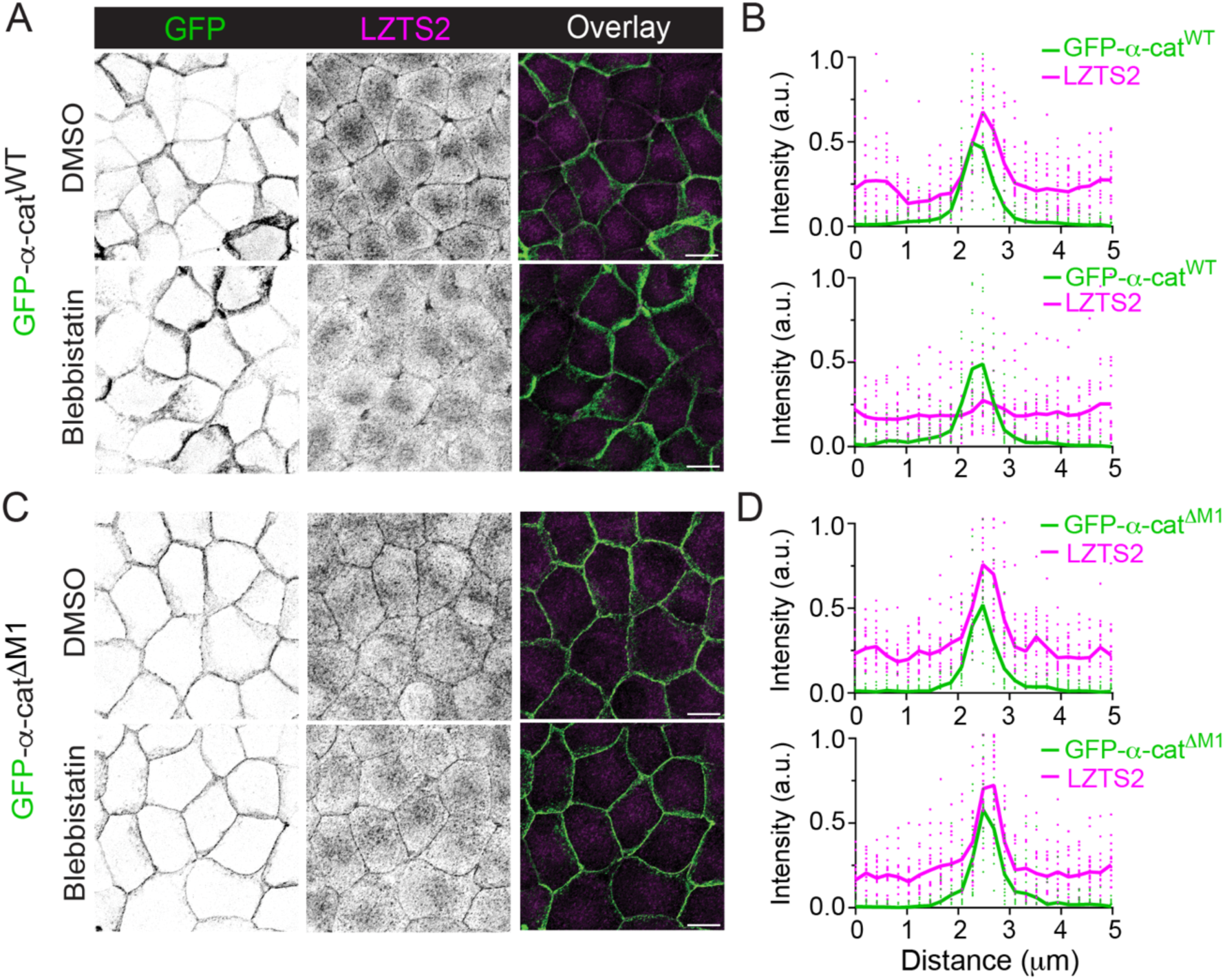
LZTS2 recruitment by α-cat 1′M1 is more uniform and refractory to myosin-inhibition compared with WT α-cat. Coverslip grown MDCK cells treated with blebbistatin (100μM, 3-hrs) reduced LZTS2 (magenta) localization to bicellular contacts in GFP-α-cat^WT^ monolayer (**A**), but not GFP-α-cat 1′M1 monolayer (**C**). Confocal *en face* views (maximum z-projections) are shown. Scale bar, 10μm. Note LZTS2 becomes more evenly distributed across bi-and tri-cellular junctions by the α-cat 1′M1 mutant WT α-cat. **(B,D)** Normalized intensity profiles of GFP-α-cat (green) and LZTS2 (magenta) based on line scans (width: 3 pixels) performed across cytosol to bicellular junctions (line scan average of 10 junctions/construct and condition).

### LZTS2 localizes to apical midbody junctions during abscission

To understand how LZTS2 depletion leads to elevated binucleation rates, we examined LZTS2 enrichment near midbody structures by immunofluorescence analysis of fixed cells, inferring early versus late stages of cytokinesis from the thickness, size and apical bias of the midbody (marked by acetylated tubulin) (Guizett et al., 2011; Hu et al., 2012; Karasmanis et al., 2019; Osswald & Morais-de-Sá, 2019). During late cytokinesis, LZTS2 could be found flanking the apically resolving midbody (Fig. 9A-C), confirming an intimate relationship between LZTS2 and the process of abscission. *En face* (x-y views) of dividing cells across different stages of cytokinesis showed progression of LZTS2 enrichment at junctions proximal to acetylated tubulin midbody structures (Fig. 9D), a pattern resembling localization of septins, anillin and non-muscle myosin II (Piekny et al., 2008; Hu et al., 2012; Karasmanis et al., 2019; Straight et al., 2005). Once the dense intercellular bridge formed, LZTS2 signal condensed into punctate junctional ring-like structures surrounding the midbody and ultimately appearing to “lasso” the midbody remnant during the final cleavage step (i.e., abscission) (Fig. 9D, far right). Progression through cytokinesis was inferred by judging acetylated tubulin thickness and shape (Fig. 9D, middle panel), since a fluorescently-tagged LZTS2 stably expressed in MDCK cells was prone to aggregation, challenging our ability to follow this process by live imaging (not shown).

**Fig. 9:**
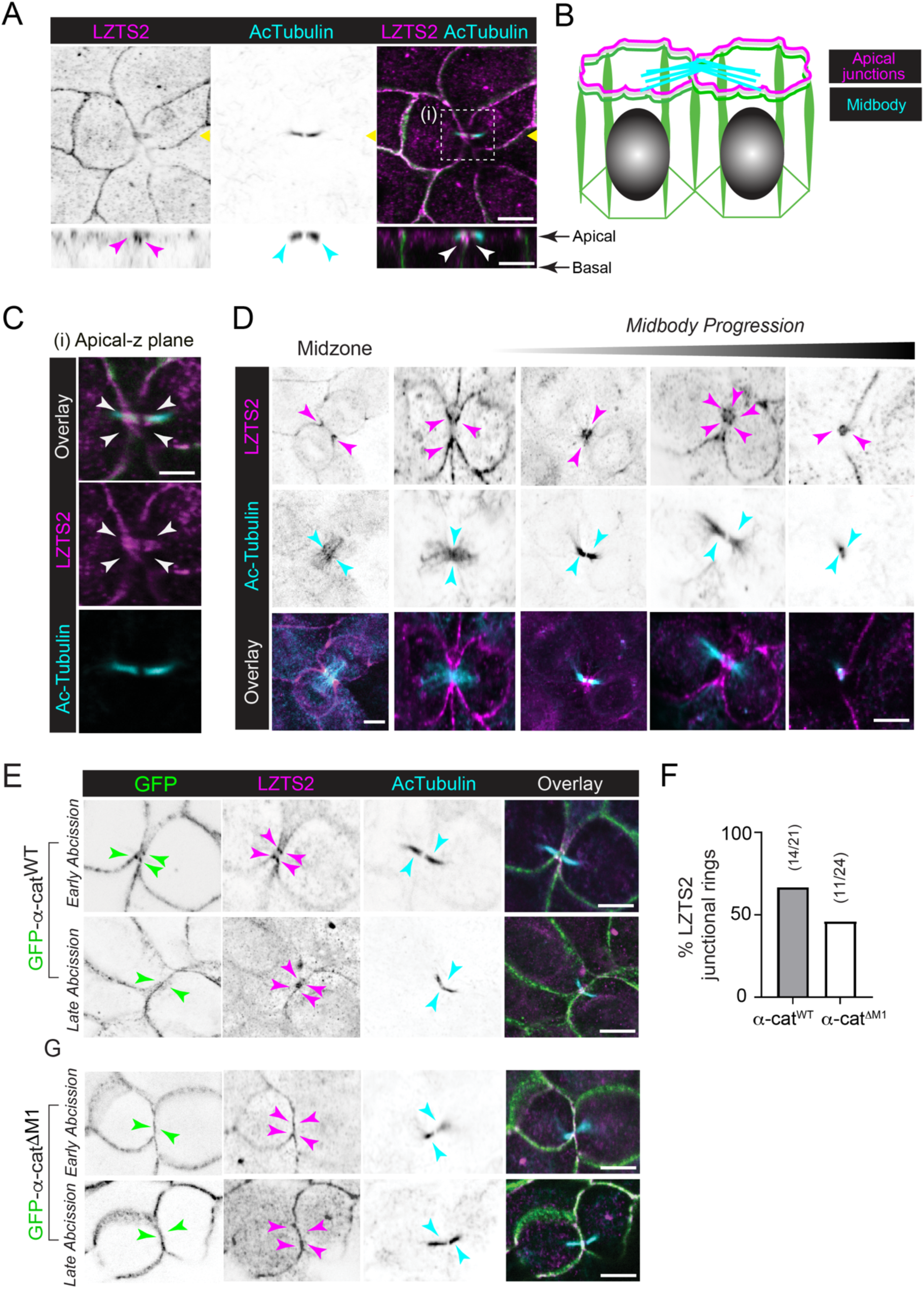
LZTS2 enriches at junctions proximal to the apically resolving midbody during abscission. **(A)** Confocal image of midbody (acetylated tubulin, cyan) flanked by LZTS2 enrichment (magenta) in α-cat^WT^ restored α-cat KO MDCK cells. Orthogonal (x-z) views shown below reveal that the midbody/abscission site localizes apically and in close proximity to LZTS2 junctional enrichment. Scale bar, 3μm. **(B)** Schematic representation of A. (**C**) Higher magnification view of midbody shown in A (white box inset, i). Arrowheads denote midbody. **(D)** Confocal images of filter grown MDCK cells fixed and immunostained for LZTS2 (magenta) and acetylated tubulin (cyan). Images capture various stages of cytokinesis and abscission. Note LZTS2 enriches at punctate junction structures that surround the midbody (note 5-point ring structure, magenta arrows). (**E-F**) Confocal images of filter-grown MDCK α-cat KO cells expressing GFP-α-cat^WT^ (upper panels) or GFP-α-cat^ι1M1^ (lower panels). Images from fixed cells captured at early versus late stages of abscission. Note LZTS2 co-localizes with GFP-α-cat^WT^ at midbody apical junctions (14 out of 21 cell pairs), whereas this structure was less frequent for 1′M1 cells (11 out of 24 cell pairs).

Since α-cat ΔM1 persistently sequesters LZTS2 to junctions, we wondered whether this mutant might have consequences for LZTS2 enrichment at the midbody. Comparison of late-stage cytokinesis images (Fig. 9E, G) revealed that 66% (14/21) of midbodies from WT α-cat-restored MDCK cells showed LZTS2 punctate ring-like enrichment, whereas fewer of these structures were seen in α-cat ΔM1-restored MDCK cells (46%,11/24) (Fig. 9F). These data suggest that persistent recruitment of LZTS2 to apical junctions may interfere with the normal recruitment pattern of LZTS2 to apical midbody junctions.

### α-cat ΔM1 shows enhanced binucleation rate over wild-type or other α-cat mutants

Evidence that α-cat ΔM1 persistently recruits LZTS2 from cytoplasmic and midbody junctions raised the possibility this mutant would impact the fidelity of cell division. We grew WT α-cat and M-domain mutant α-cat-restored MDCK cells on filters for 14 days and found that the α-cat 1′M1 mutant was associated with the highest rate of binucleation (Fig. 10A-C), even compared with other α-cat M-domain and FABD+ mutants (Fig. 1, 3 and 4). Thus, these data show that an α-cat mutant that constitutively exposes M2-M3 helices and sequesters LZTS2 from the midbody is associated with binucleation, likely through a defect in cytokinesis. Importantly, the ability of α-cat ΔM1 to promote binucleation is also seen when expressed in parental MDCK cells (Fig. S5), suggesting this mutant can function as a dominant inhibitor of cytokinesis, likely through sequestration of a factor required for this process (i.e., LZTS2). Persistent recruitment of vinculin driven by the α-cat1′M2,3 mutant (through exposing M1 helices for vinculin binding) did not appear to drive binucleation, although this mutant does appear to impact mitotic rounding (Fig. S6), consistent with previous study (Monster et al., 2020). Collectively, these data suggest that α-cat mechanosensitivity is normally tuned to allow faithful cytokinesis and abscission, where enhanced mechano-activation of α-cat’s M-domain might perturb this process through persistent sequestration of the abscission factor, LZTS2.

**Fig. 10:**
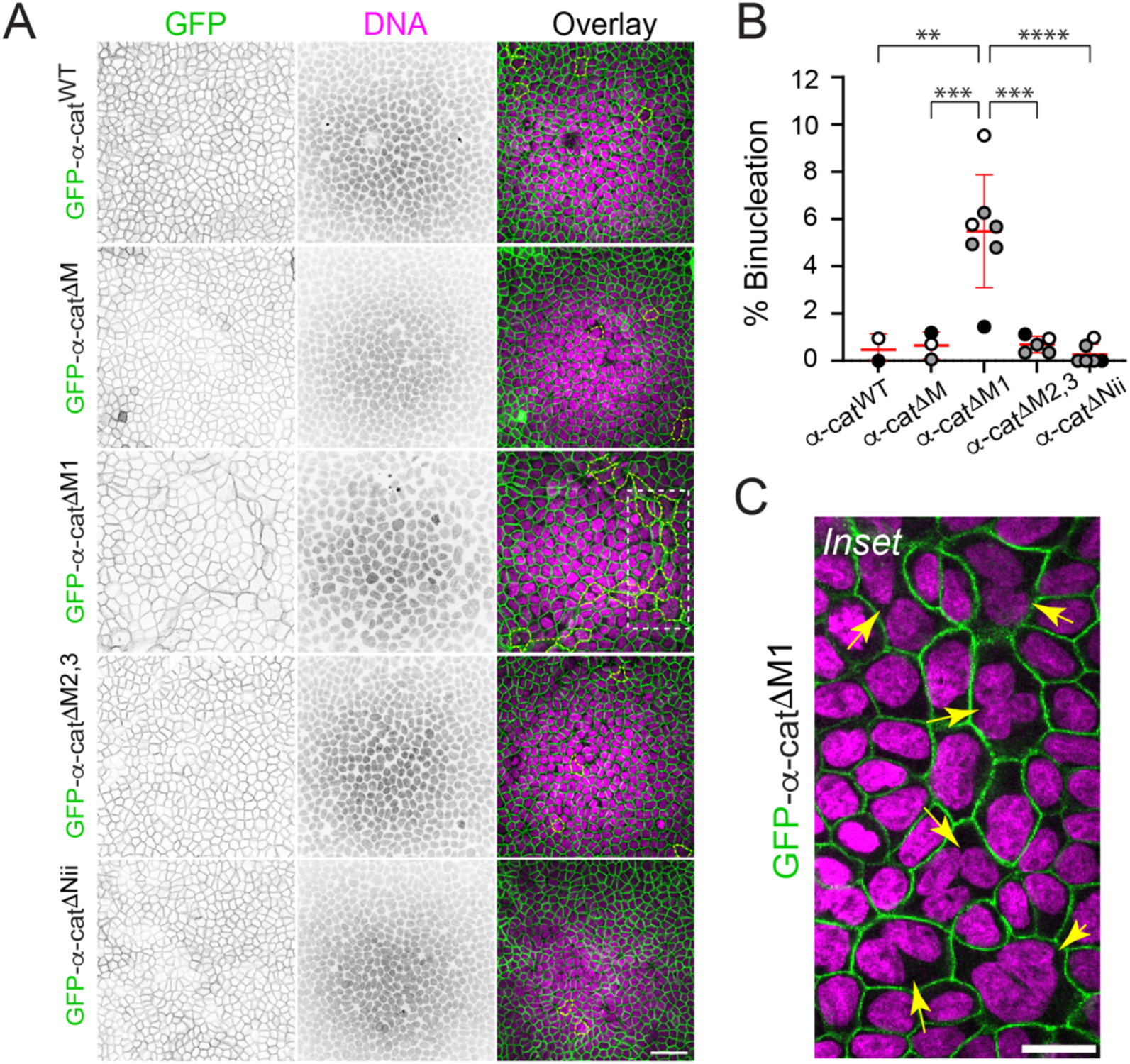
α-cat-1′M1 monolayers show enhanced binucleation rate over wild-type and other α-cat mutants. **(A)** Confocal *en face* view (maximum z-projection; grayscale inverted) images showing GFP-α-cat^ι1M1^ expressing cells exhibit polyploidy after 12 days grown on filters. GFP-α-cat (green) and LZTS2 (magenta), multinucleated cells (outlined, yellow dashed line). Representative images from four independent experiments are shown. Scale bar, 40μm. **(B)** Quantification of multinucleation rate in different GFP-α-cat constructs expressing MDCK α-cat KO cells. Two-tailed unpaired t test, **p<0.01, ***p<0.001, ****p<0.0001. Graph indicates mean ± SD. Data from three independent experiments. **(C)** The rectangular regions framed in white in (A) are enlarged to show detail. Scale bar, 20μm. Yellow arrows denote bi-or multinucleated cells.

**Fig. 11:**
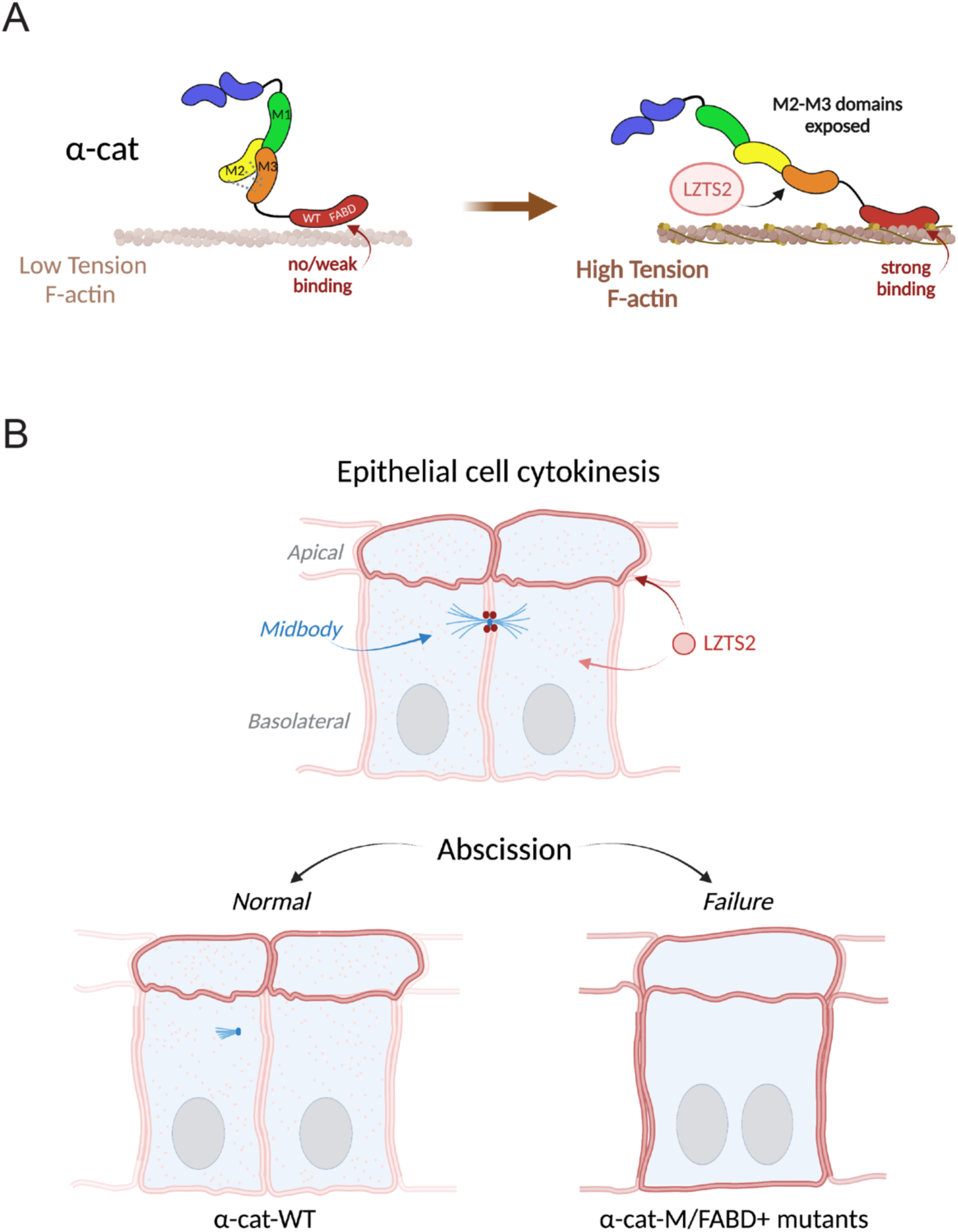
Model. **(A)** α-cat M2-M3 domain unfurling traps LZTS2. **(B)** Persistent α-cat M-domain opening sequesters LZTS2 to cellular membranes, limiting access to apical midbody-junctions during late-stage cytokinesis.

## DISCUSSION

Studies suggest that an abortive cytokinesis program can serve as means to generate multinuclear polyploid cells across a range of cell-types and conditions, particularly where extra genome copies may be advantageous (Bailey et al., 2021; Øvrebø & Edgar, 2018; Schoenfelder & Fox, 2015). For example, cardiomyocytes become binucleated through a developmentally timed defect in cytokinetic furrow progression (Engel et al., 2006; Normand & King, 2010; Soonpaa et al., 1996); macrophages multinucleate downstream of bacteria/Toll-Like Receptor signals that antagonize cytokinesis (Herrtwich et al., 2016). Even simple barrier epithelia become binucleated after mechanical injury due to failed cytokinesis (Cao et al., 2017b; Lazzeri et al., 2019; Weng et al., 2022), where increased DNA copies lead to larger cells that manifest less junctional surface relative to epithelial area coverage resulting in reduced barrier leak (Cohen et al., 2018; Peterson & Fox, 2021; Schoenfelder & Fox, 2015). Collectively, these observations suggest abortive cytokinesis programs may be a design feature of the mitotic division to favor various adaptive outcomes. Mechanistically, excessive integrin-extracellular matrix adhesive tension along epithelial wound fronts appears to be one cause of cytokinesis failure, where exuberant stress fiber formation can interfere with contractility of the cytokinetic ring leading to division failure (Uroz et al., 2018, 2019). Whether other mechanosensitive players similarly interfere with cytokinesis has remained unclear.

In this study, we take advantage of rationally designed mutant forms of α-cat to implicate adherens junction mechanosensitivity as a possible upstream mediator of cytokinetic fidelity. We find that a force-desensitized α-cat actin-binding mutant (α-cat-H0-FABD^+^), which shows enhanced binding to F-actin *in vitro* (Ishiyama et al., 2018) and stronger epithelial sheet integrity through enhanced coupling to lower tension cortical actin networks (Wood et al., 2023), can interfere with cytokinesis. Since this interference is not observed in single cells or cells lacking cadherins (Fig. S1), the α-cat-H0-FABD^+^ mutant likely promotes cytokinesis failure through cadherin-catenin junction complexes engaged in cell-cell adhesion (Le Bras & Le Borgne, 2014), rather than exclusive extra-junctional homodimer functions. In addition, this interference is not mediated by the α-cat ABD alone but requires both M-and H0-FABD^+^ domains to drive cytokinesis failure. Thus, a normal functioning α-cat is required for cytokinesis fidelity in epithelial monolayers.

Evidence the α-cat M-domain contributes to cytokinetic interference aligns with recent evidence that heterozygous missense variants in *CTNNA1*/α-cat (M-domain) cause a form of macular dystrophy through driving an age-associated multinucleated epithelial phenotype (Saksens et al., 2016). Butterfly-shaped Pigmentary macular Dystrophy (BPD) is an autosomal dominant eye disease characterized by bilateral accumulation of pigment in the macular area that resembles the wings of a butterfly. Remarkably, a forward genetic screen for similar eye defects in mice independently identified an α-cat M-domain missense mutation L436P, where *Ctnna1*^tvrm5^ ^(L436P)^ mice reveal earliest stages of disease pathogenesis may initiate through progressive multinucleation of retinal pigment epithelial cells. While this phenotype was stronger in mice homozygous for the *Ctnna1*^tvrm5^ ^(L426P)^ mutation, RPE defects were also seen in heterozygotes, suggesting a gene-dosage effect. It is important to note that cell-fusion is also a means to generate multi-nuclear epithelial cells (Bailey et al., 2021; Øvrebø & Edgar, 2018; Gjelsvik et al., 2019), where fly L426P mutant α-cat appears to enhance this fusion phenotype after injury (Dehn et al., 2021). However, expression of BPD missense variants in MDCK is associated with an elevated binucleation rate via cytokinesis failure (Fig.4, Fig. S2), suggesting different paths to multi-nucleation across species. How α-cat mutants generate junction signals that serve to sustain the polyploid state will require further investigation.

There are many routes to cytokinesis failure, the best-known involving reduced or elevated Rho-signaling (Kamijo et al., 2006; Konstantinidis et al., 2015; Lordier et al., 2008; van de Ven et al., 2016; Zhou et al., 2013). Curiously, α-cat mutants that interfere with M- or FABD mechanosensitive functions lead to cytokinesis failures during the later stages of this process. This suggests α-cat may interfere with abscission, which in polarized epithelia is apically biased and must resolve in close proximity with apical junctions (Osswald & Morais-de-Sá, 2019; Ott, 2016; Tassan et al., 2017; Higashi and Miller, 2017; Bai et al., 2019). Consistent with this model, we show that an α-cat M-domain salt-bridge mutant designed to mimic a constitutively unfurled state (i.e., force-independent R551A mutant (Barrick et al., 2018) can recruit a previously suggested “abscission factor”, Leucine Zipper Tumor Suppressor 2 (LZTS2) (Johnson et al., 2013; Maru, 2009; Sudo & Maru, 2007), where persistent mechanosensitive recruitment of LZTS2 to AJs may sequester LZTS2 from the midbody and its participation in abscission (Maru, 2009; Sudo & Maru, 2007). While we and others find LZTS2 depletion leads to binucleation, molecular details remain unclear. LZTS2 is a poorly understood scaffold protein previously localized to centrosomes and midbody (Sudo & Maru, 2007; Sudo and Maru, 2008). While there is some evidence LZTS2 is functionally linked to microtubule severing (Sudo and Maru, 2008; Maru, 2009), recent proteomics screens also place LZTS2 within a protein network centered around Afadin, another mechanosensitive binding partner of α-cat (Baskaran et al., 2021). Future work is required to understand how this protein network dynamically interacts with mechanosensitive cadherin-catenin adhesions to guide cytokinesis to completion. Evidence that α-cat mechanosensitive mutants can interfere with cytokinesis, reinforces the idea that midbody abscission critically depends on a tension release step (Andrade et al., 2022; Andrade & Echard, 2022; Hatte et al., 2017; Herbomel et al., 2017; Lafaurie-Janvore et al., 2013; Rabie et al., 2021).

## STUDY LIMITATIONS

Live imaging MDCK cells during cytokinesis, particularly the abscission step which resolves apically in epithelia, is challenging for a system where the timing and location of division events are not predictable across MDCK cell cultures. Thus, future work on the contribution of α-cat mechanosensitivity to cell division will greatly benefit from a system where cells are large, events are abundant, and the plane of division is uniformly oriented (e.g., as in cleaving Xenopus embryos).

It is also important to disclose that the propensity of α-cat M- and FABD+ mutants to interfere with cytokinesis may be due to features of the MDCK α-cat KO/reconstitution system. Indeed, α-cat CRISPR KO MDCK clone 2.2 (and other clones) show high levels of binucleation in up to 20% of MDCK cells (Fig. S7). This finding was originally missed, because we largely focused attention on GFP-α-cat WT and α-cat-mutant “rescue” MDCK cells. While we presume this binucleation is due to cytokinesis failure, single cell imaging of α-cat KO cells led to few captured events due to their reduced spreading capacity during live imaging and loss during mitotic rounding (Fig. S1). Since GFP-α-cat WT and α-cat-mutant “rescued” α-cat CRISPR KO MDCK cells show lower binucleation rates (2% - 6%; Figs 1 and 10), these data suggest normal α-cat mechanosensitivity appears required for cytokinetic fidelity, where *either* loss of α-cat or altered α-cat M-domain function can interfere with this process.

## METHODS

### Lentiviral plasmid generation and constructs

All GFP-α-cat constructs were synthesized by Vectorbuilder using the αE-catenin human CTNNA1 sequence. The ability of eGFP to spontaneously dimerize and potentially impact α-cat functions was disrupted by incorporating an A206K mutation (Zacharias et al., 2002)). All constructs listed in the Key Resources table will be submitted to Addgene with detailed vector maps per NIH guidelines.

### Cell culture and stable cell line selection

MDCK II cells were maintained in Dulbecco’s Modified Eagle’s Medium (DMEM, Corning), containing 10% fetal bovine serum (FBS, Atlanta Biologicals or JRS Scientific), 100 U/ml penicillin and 100 μg/ml streptomycin (Corning). α-cat/*Ctnna1* knockout MDCK cells were generated using CRISPR-Cas9 system as described in Wood et al., 2023 BioRxiv. Briefly, RNA guides were designed targeting α-cat sequences in exons 2 and 4. Guide RNA (gRNA) targeting different canine *CTNNA1* exons were designed with CHOPCHOP online tools.^45^ Sequences of oligonucleotides were as follows: 5’-GAAAATGACTTCTGTCCACACAGG -3’ (exon 2, covering the MTSVHTG protein sequence) and 5’-AGTCTAGAGATCCGAACTCTGG - 3’ (exon 2, covering the SLEIRTLA sequence). Single guide RNA (sgRNA), Cas9 nuclease (HiFi) and duplex buffer were purchased from Integrated DNA Technologies. RNAs were reconstituted and diluted to 5 μM with duplex buffer; Cas9 protein to 10 μM with PBS. MDCK cells were plated in 12-well plates at a seeding density of 1.0 x 10^5^ cells in 1 mL DMEM-complete a day before. For one reaction, sgRNA (20 μL), Cas9 (15 μL), DMEM (30 μL, no serum and antibiotics) and Lipofectamine RNAiMax (4 μL, Invitrogen) were mixed and incubated at room temperature for 20 min. The complex was treated to the cells in complete medium and incubated overnight at 37 °C and 5% CO_2_. The medium was changed and the cells were allowed to recover for one day. Cells were split to maintain 30-50% confluency. The sgRNA-Cas9 treatment was repeated 3 times. Cells were expanded and sorted with a flow cytometer (FACSMelody, BD) to 96-well plates to grow single cell colonies. The colonies were screened for low (< 5 ng/well at confluent) α-cat expression using a α-cat C-terminal antibodies. The selected colonies were verified with western blot. Knockout cell line (2.2 clone) was chosen based on lowest level of α-cat isoform lacking N-terminal sequences. α-cat^KO2.2^ MDCK were restored with wild-type and mutant α-cat forms by lentiviral infection, selected in puromycin (xug/uL). GFP-α-cat-positive cells were flow sorted to ensure even expression across constructs.

### Antibodies

The following primary antibodies were used: polyclonal rabbit anti-α-cat (C3236, Cell Signaling), hybridoma mouse anti-α-catenin (5B11, (Daugherty et al., 2014)), rabbit anti-GAPDH (Cat# sc-25778, Santa Cruz, now discontinued), polyclonal rabbit anti-GFP (A11122, Invitrogen) and Phalloidin-488 or -568 (A12379, Invitrogen). Secondary antibodies for Western blotting included HRP-conjugated goat anti-mouse and -rabbit antibodies (Bio-Rad), or fluorescently labeled donkey anti-mouse and -rabbit antibodies (680RD or 800RD, LiCor Biosciences). Secondary antibodies for immunofluorescence included IgG Alexa Fluor 488 or 568-conjugated goat anti-mouse or -rabbit antibodies (Invitrogen).

### Immunofluorescence and Imaging

Cells were grown on cell culture inserts (Falcon), fixed in 4% paraformaldehyde (Electron Microscopy Services, Hatfield, PA) for 15’, quenched with glycine, permeabilized with 0.3% Triton X-100 (Sigma), and blocked with normal goat serum (Sigma). Primary and secondary antibody incubations were performed at RT for 1h, interspaced by multiple washes in PBS, and followed by mounting coverslips in ProLong Gold fixative (Life Technologies). Images of α-cat & F-actin localizations were captured with Nikon A1R Confocal Laser Point Scanning microscope using NIS Elements software (Nikon) with GaAsP detectors and equipped with 95B prime Photometrics camera, Plan-Apochromat 60x/1.4 objective. Confocal Z-stacks were taken at step size of 0.3μm.

### Scratch wound assay

For Fig. 1F-H, Fig. 4C-D, MDCK cells were plated for 10days on 12-well Falcon cell culture inserts (BD Falcon 353494; high-density 0.4μm), wounded with vacuum suction pipette, rinsed with PBS, and recovered in complete DMEM media for 40hrs before fixation & immunostaining (see immunofluorescence and imaging section for details). For Fig. S2B-C, cells were imaged on Nikon Biostation IM-Q with the slide holder module as previously described (Ishiyama et al., 2018). Briefly, MDCK cells were plated for 24hr on LabTek #1 4-well chamber slide (43300-776, Thermo Fisher Scientific), wounded with vacuum suction P200 micropipette tips and recovered for 2hrs. Prior to imaging, complete DMEM media was replaced with FluoroBright DMEM (Life Technologies). Cells were imaged with 20X objective every 10min (both phase contrast and fluorescent channels) on Biostation IM-Q at 37C, 5%CO2, for 16hrs. 4 fields of view (FOV) were captured along the wound edge. Instrument controlled by Biostation IM software, version 2.21 build 144.

### Biotin ligase (BioID) screen for α-cat M-domain proximity partners

BirA*-α-cat fusion proteins were generated using wild-type α-cat and the R551A salt-bridge mutant previously established to enhance a-cat accessibility to tension-sensitive partners, such as vinculin (Ishiyama et al., 2013). Plasmids were expressed in HEK293 T-REx cells and subjected to LC-MS/MS. Since WT α-cat recovered more spectra than the α-cat R551A mutant, we applied a total spectral count-based normalization. We forced only proteins detected with a SAINT FDR ≤1% in one of the conditions to be visualized, and only those proteins detected with at least 5 spectra in 1 condition are displayed.

Key resources Table.

| REAGENT or RESOURCE | SOURCE | IDENTIFIER |
| --- | --- | --- |
| **Antibodies** |  |  |
| Anti-α-catenin | Cell Signaling | Cat#3236 |
| Anti-α-catenin (5B11) | | https://www.pnas.org/doi/10.1073/pnas.1308663111 |
| LZTS2 rabbit polyclonal antibody | Proteintech | 15677-1-AP |
| Acetylated-tubulin (6-11B-1) mouse monoclonal | Sigma-Aldrich | T7451 |
| Vinculin (mouse monoclonal) | Sigma-Aldrich | V9131 |
| Phalloidin-Far Red | Invitrogen | A22287 |
| GAPDH | Santa Cruz | Cat# sc-25778 |
| Anti-GFP | Invitrogen | A11122 |
| Alexa Fluor 568 phalloidin | Invitrogen | Cat#A12380 |
| Alexa Fluor 488 phalloidin | Invitrogen | Cat#A12379 |
| Goat anti-rabbit Alexa Fluor 488 | Invitrogen | Cat#A11008 |
| Goat anti-mouse Alexa Fluor 568 | Invitrogen | Cat#A11004 |
| HRP-conjugated goat anti-mouse | BioRad | Cat#1706516 |
| HRP-conjugated goat anti-rabbit | BioRad | Cat#1706515 |
| Donkey anti-mouse | LiCor Biosciences | P/N 925-68072 |
| Donkey anti-rabbit | LiCor Biosciences | P/N: 926-32213 |
| **Chemicals, Peptides, and Recombinant Proteins** |  |  |
| Hoechst 33342 | ThermoFisher Scientific | Cat#62249 |
| ProLong Gold Antifade Mountant | ThermoFisher Scientific | Cat#P36934 |
| **Experimental Models: Organisms/Strains** |  |  |
| Madin-Darby canine kidney II cells | Gumbiner Lab (1996) | Heidelberg strain (Kai Simmons Lab) |
| A431D cells | Trojanovsky Lab | E-cad negative variant of A431 epidermoid skin cancer |
| **Software and Algorithms** |  |  |
| FIJI/ImageJ (version: 2.1.0/1.53c) | Schneider, CA et al., 2012 | https://imagej.nih.gov/ij/ |
| Graphpad Prism |  | https://www.graphpad.com/scientific-software/prism/ |
| **Other** |  |  |
| BioStation Slide Holder | Nikon Instruments |  |
| Axioplan2 epifluorescence microscope | Zeiss; 20x objective (Air) | AxioCAM HR Camera with AxioVision 4.8 software |
| A1R Confocal | Nikon Instruments | Laser Point Scanning, GaAsP detectors, Plan-Apochromat 60x/1.4 objective |
| **Plasmids** |  |  |
| LV-monomeric EGFP-αE-catenin (WT, missense, M-domain and FABD mutants) | Gottardi Lab | Vectorbuilder; Will submit to Addgene with vector maps |
| LV-mEGFP:hCTNNA1_Wild-type |  |  |
| LV-mEGFP:hCTNNA1_H0-FABD+ | (RAIM<sup>669-672</sup>>GSGS) |  |
| LV-mEGFP:hCTNNA1_ΔM-domain | Δ273-635 |  |
| LV-mEGFP:hCTNNA1_ΔM1-domain | Δ271-396 |  |
| LV-mEGFP:hCTNNA1_ΔM2,3-domain | Δ397-509 |  |
| LV-mEGFP:hCTNNA1_ΔNii domain |  |  |
| LV-mEGFP:hCTNNA1_R551A |  | M2-M3 salt-bridge disrupting mutant |
| LV-mEGFP:hCTNNA1_L436P |  | *Ctnna1* eye phenotype via forward genetic screen |
| LV-mEGFP:hCTNNA1_E307K |  | CTNNA1 butterfly eye mutant |

## Author Contributions

Conceptualization: MI and CJG; Experimentation/Analysis: YW, AY, CG SK, JMQ, ASF, SL; Graphics: PML; Writing: YW and CJG; Editing: YW, CJG and ASF; Supervision: CJF Funding: MI, AC, CJG

## Condensed title

α-catenin and cytokinesis fidelity

## ACKNOWLEDGEMENTS

This work relied on the following Northwestern University services and core facilities: Center for Advanced Microscopy (NCI CCSG P30 CA060553, NCRR 1S10 RR031680, 1S10OD021704) and Flow Cytometry (NCI CA060553, 1S10OD011996, 1S10OD026814).

## FIGURES AND LEGENDS

**Fig. S1:**
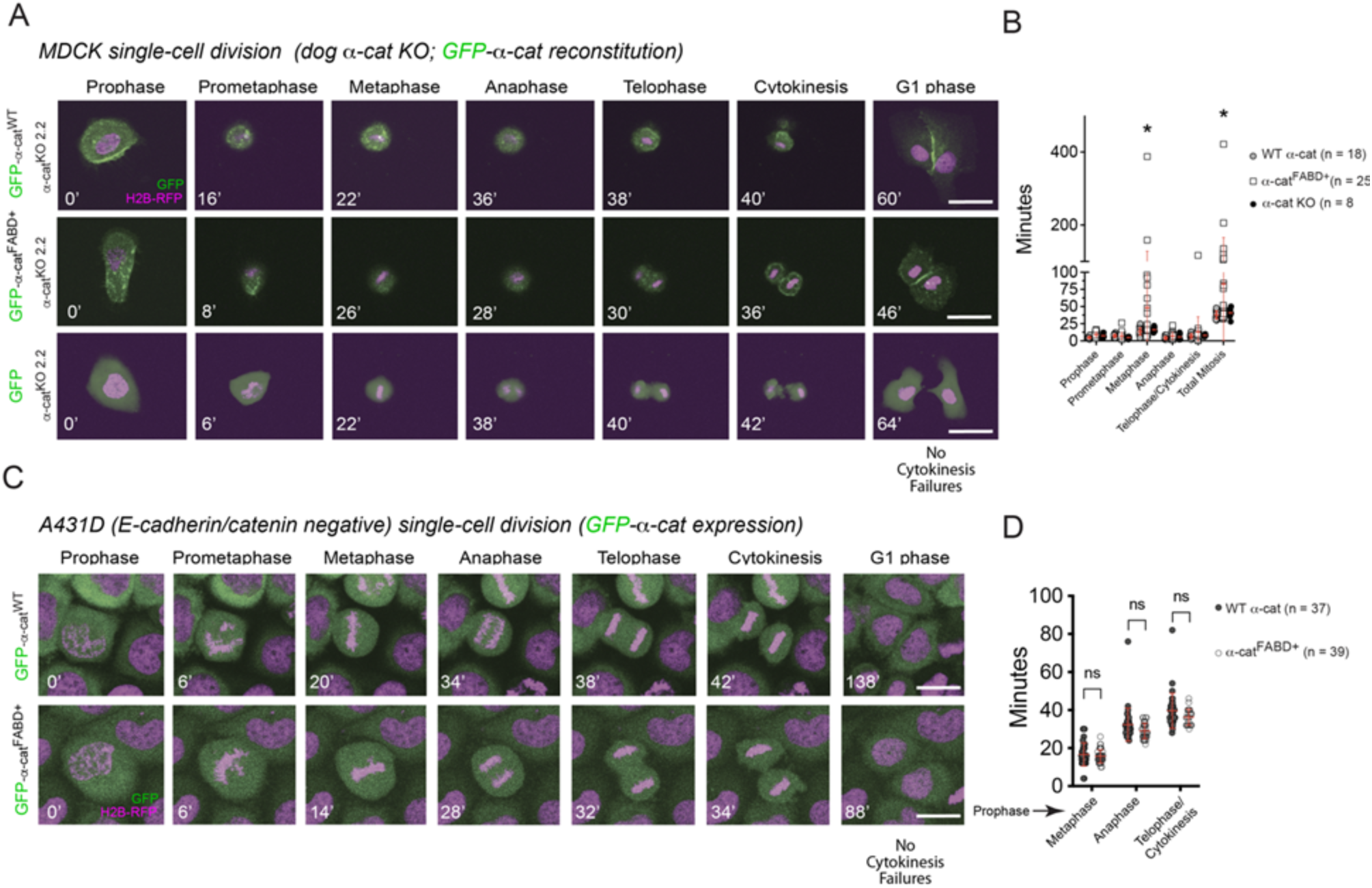
α-cat^H0-FABD+^ -induced cytokinesis failure does not occur in single cells. **(A)** Live imaging analysis of MDCK α-cat KO cells restored with α-cat^WT^ (Top), α-cat^H0-FABD+^ (Middle) or with GFP alone (Bottom) seeded at low density to capture single-cell mitosis events. Metaphase, anaphase, telophase and cytokinesis phases shown. Time stamp in minutes (‘) shown in white; α-cat-GFP (green), nuclei (magenta). Scale bar, 50μm **(B)** Graph quantification of mitosis phase lengths shown. While α-cat^H0-FABD+^ single cells spend longer in metaphase (*p < 0.05 by unpaired t-test), no cytokinesis failures were observed. Comparatively few cell divisions were captured for α-cat KO cells (N = 8 versus 18 (α-cat^WT^) and 25 (α-cat^H0-FABD+^), due to their inability to adhere during imaging. **(C)** Live imaging analysis of pan-cadherin-negative A431D cells (Lewis et al., 1997) restored with α-cat^WT^ (Top) or α-cat^H0-FABD+^ (Bottom). Metaphase, anaphase, telophase and cytokinesis phases shown. Time stamp in minutes (‘) shown in white; α-cat-GFP (green), nuclei (magenta). Scale bar, 30μm **(D)** Graph shows mitotic phase lengths starting from prophase, where N = 37 and 39 for α-cat^WT^ or α-cat^H0-FABD+^, respectively.

**Fig. S2:**
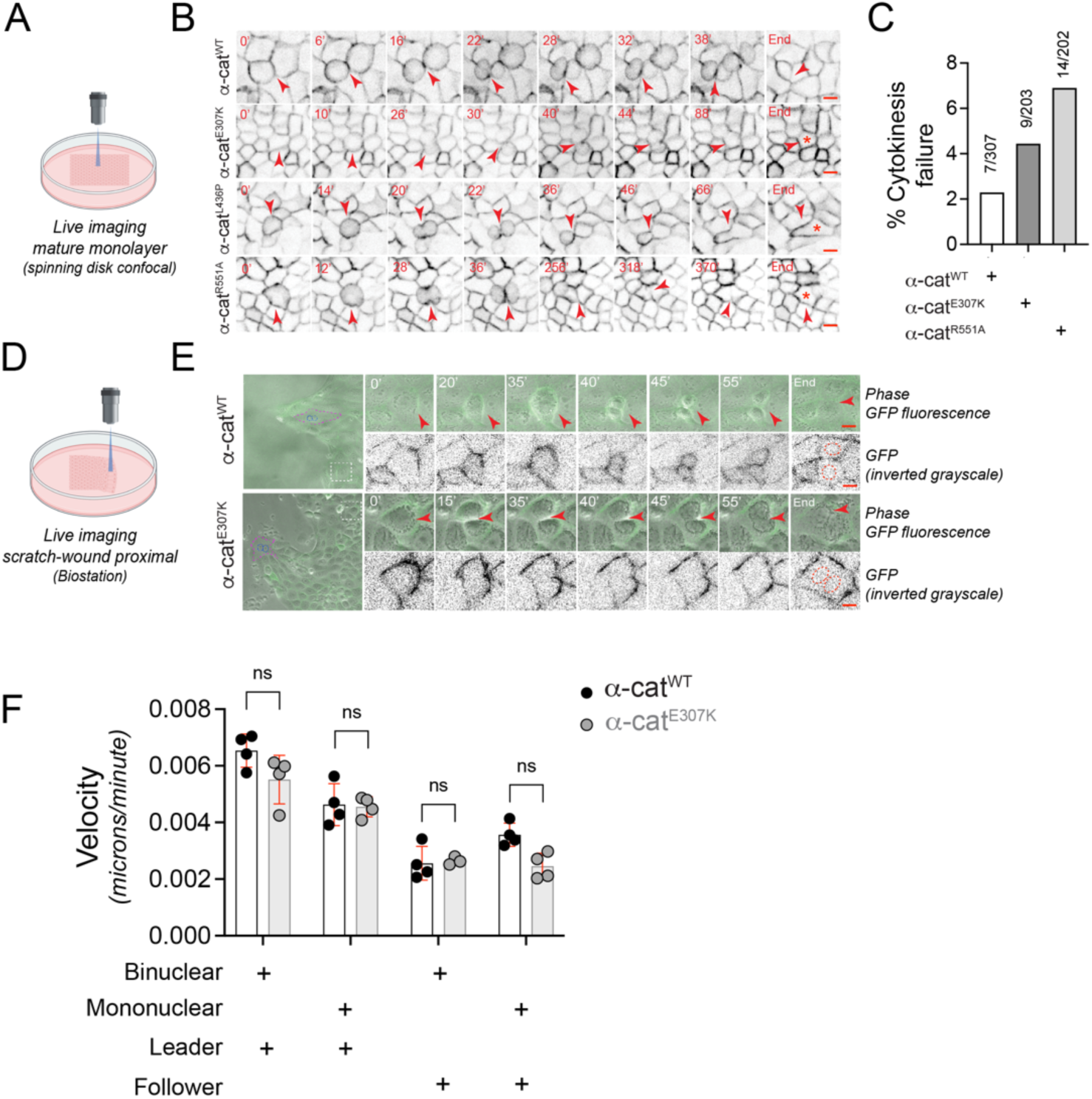
α-cat M-domain missense mutants induce failed cytokinesis. **(A-B)** Live imaging analysis of MDCK α-cat KO cells restored with α-cat^WT^, α-cat^E307K^ and α-cat^R551A^ and during mitosis phases (α-cat-GFP, black-inverted grayscale). Time stamp in minutes (‘,red). Red arrowheads show dividing cells; asterisk shows frame where cytokinesis fails. Scale bar, 10μm. (**C**) Graph shows elevated rate of cytokinesis failures in MDCK α-cat KO cells expressing GFP-α-cat^E307K^ and GFP-α-cat^R551A^ compared with GFP-α-cat^WT^ **(D-E)** Time-lapse images showing evidence of cytokinesis failure at MDCK wound fronts (Biostation). **(F)** Graph shows no clear differences in migration speeds of α-cat^E307K^ mutant versus α-cat^WT^ cells by unpaired t-test, although binucleation is associated with faster migration speeds. Analysis of the α-cat^E307K^ mutant was chosen as a test-case to rule-in or -out a model where enhanced binucleation observed at wound fronts (Fig. 4) is due to enhanced migration speeds of binucleated α-cat mutant cells. Data in F do not strongly support this idea. Other α-cat mutants (α-cat^R551A^, α-cat^L436P^) were not examined.

**Fig. S3:**
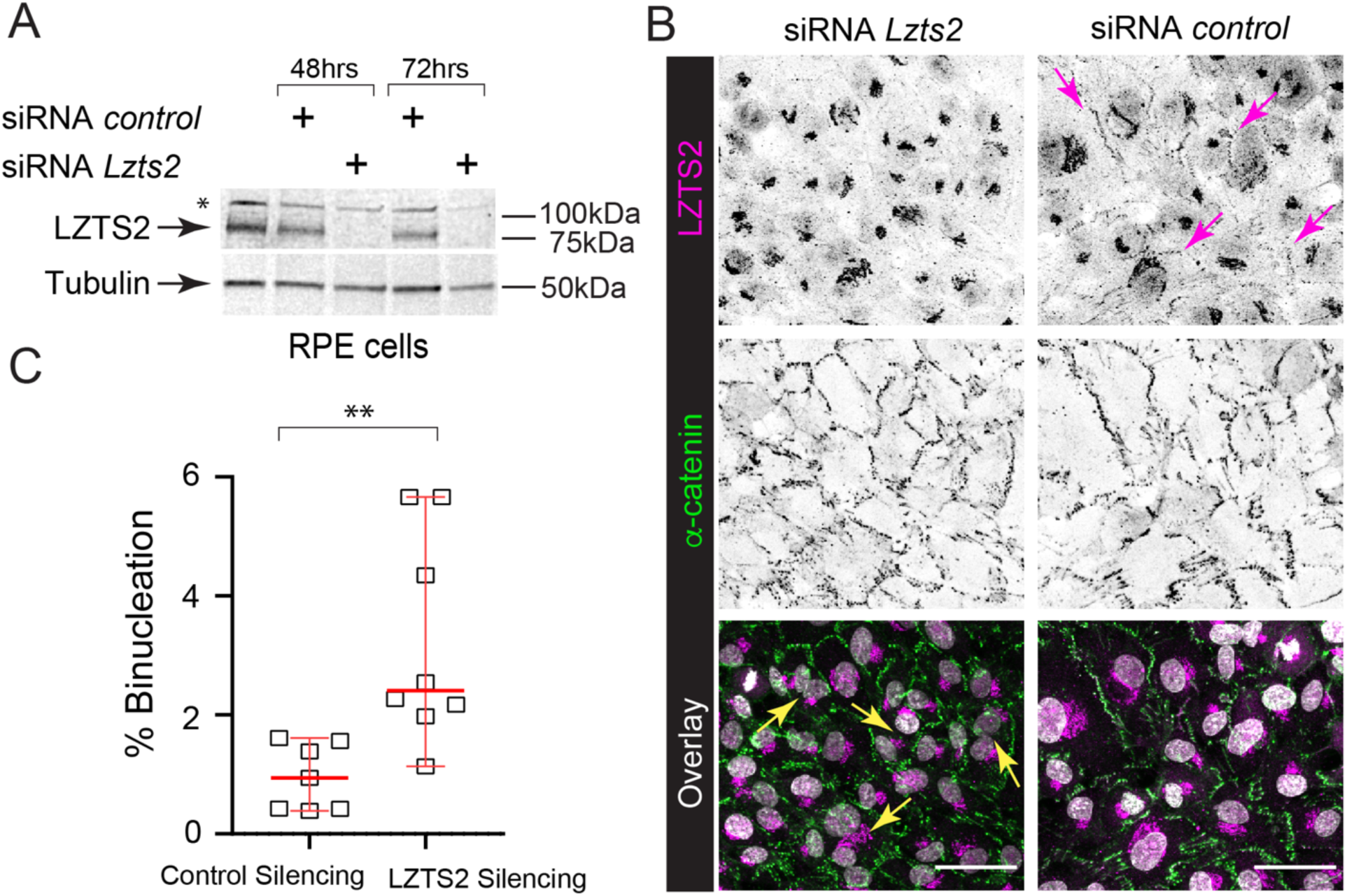
LZTS2 knock-down in RPE cells leads to binucleation. **(A)** Immunoblots of total cell extracts showing efficient transient knockdown of LZTS2 expression using the mixture of two different LZTS2 siRNAs. Asterisk (*) denotes non-specific band refractory to knock-down and possibly contributing to Golgi localization. **(B)** Confocal *en face* view (maximum z-projection; inverted grayscale) showing efficient transient knockdown (48hrs) of LZTS2 at cell-cell contacts (denoted by magenta arrows) using the mixture of two different LZTS2 siRNAs. LZTS2-deficient RPE cells exhibit binucleation (denoted by yellow arrows). LZTS2 (magenta), α-cat (green), nuclei (gray). Scale bar, 50μm. **(C)** Percentage of binucleated RPE cells upon transient knockdown of LZTS2 expression using the mixture of two different LZTS2 siRNAs. Two-tailed unpaired t test, **p<0.01. Graph indicates mean ± SD. Data from one experiment.

**Fig. S4:**
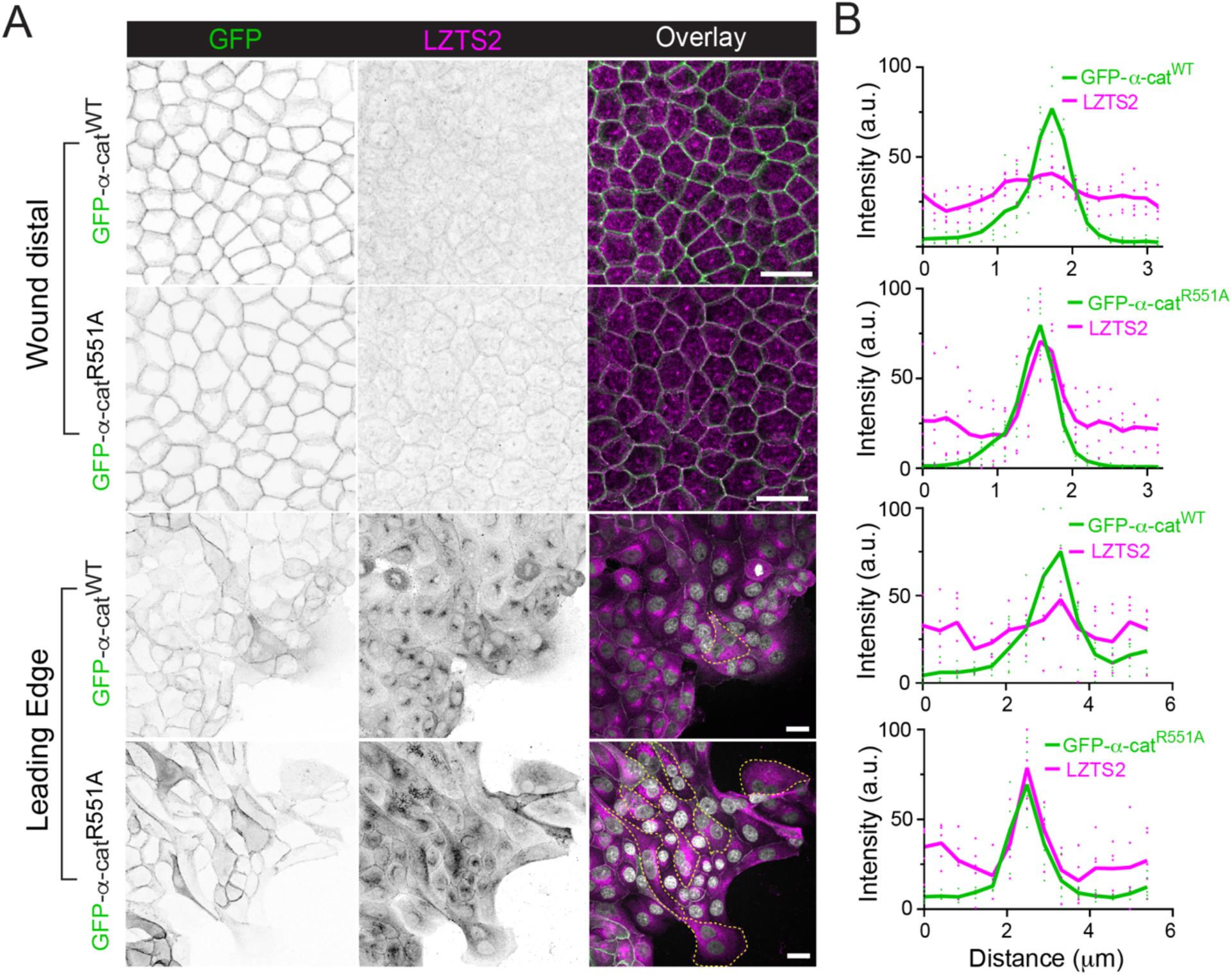
α-cat M-domain R551A salt-bridge mutant only modestly sequesters LZTS2 to apical junctions relative to wild-type α-cat. **A)** Confocal *en face* views (maximum z-projection; grayscale inverted) of α-cat^WT^ and α-cat^R551A^ restored α-cat KO MDCK cells distal and proximal to the wound front (leading edge). Cells expressing α-cat^R551A^ modestly enriches LZTS2 at cellular junctions. α-cat (green), LZTS2 (magenta), Scale bar, 10μm. Representative images from four independent experiments are shown. Yellow dashed outlines indicate binucleated cells. **(B)** Normalized intensity profiles of GFP-α-cat (green) and LZTS2 (magenta) based on line scans (width: 3 pixels) performed across cytosol to bicellular junctions (line scan average of 5 junctions/construct).

**Fig. S5:**
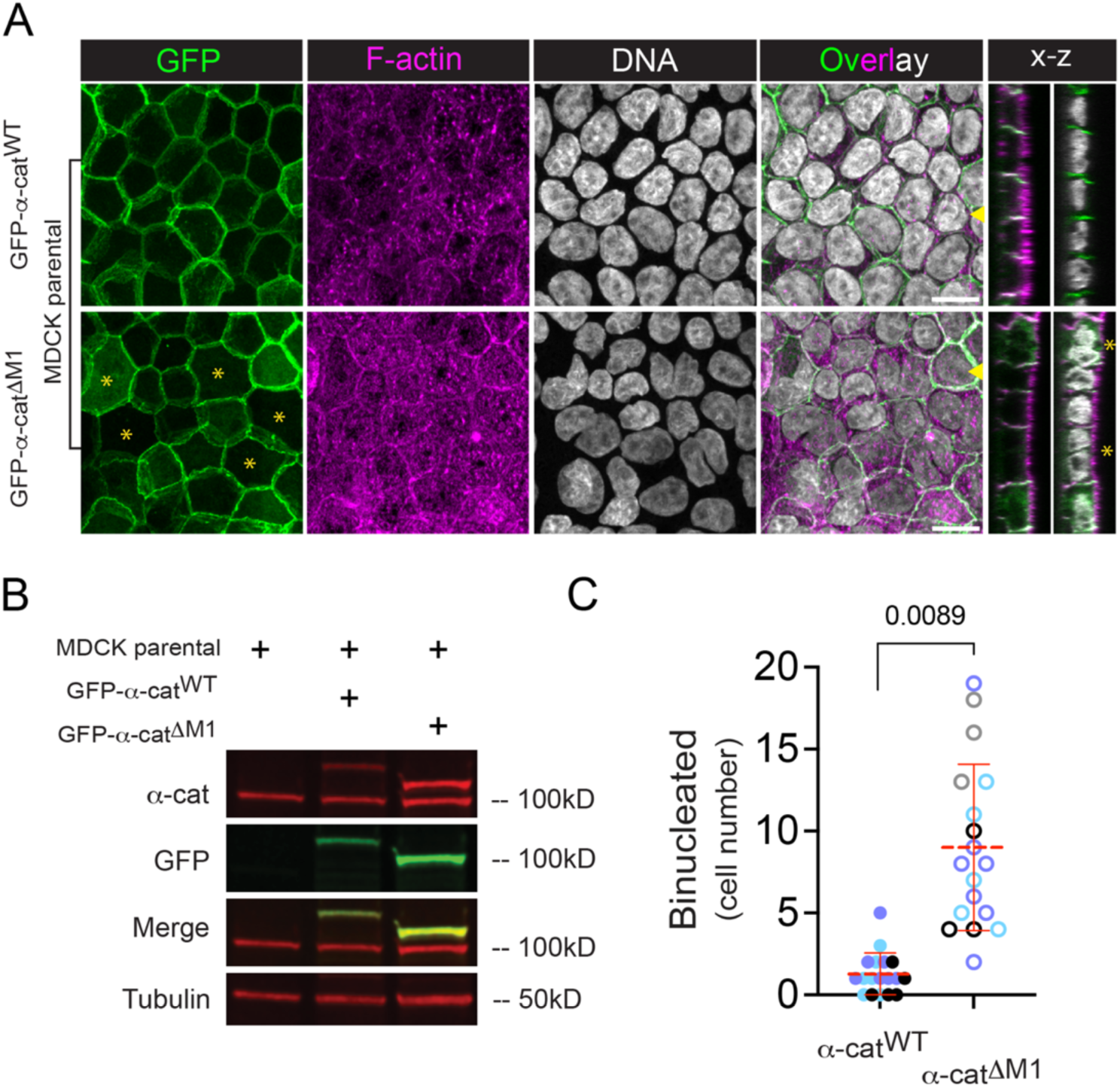
α-cat-1′M1 is sufficient to induce binucleation in parental MDCK cells. **(A)** Confocal images of filter-grown parental MDCK cells stably expressing GFP-α-cat WT or GFP-α-cat1′M1 mutants. α-cat/GFP (green), F-actin (magenta) and DNA (gray) reveal abundant binucleation associated with α-cat1′M1 expression (yellow asterisks). Maximum z-projection is shown; x-z stacks of shown to right, where yellow arrowheads mark position of optical slice). Scale bar 10μm. Representative images from multiple independent experiments are shown. **(B)** Immunoblot of GFP-α-cat construct expression in MDCK parental cells with tubulin loading control. (**C**) Binucleation rate counted manually across 18 fields of view, where distinct symbol colors reflect different biological replicates. Statistical significance by t-test (p = 0.0089).

**Fig. S6:**
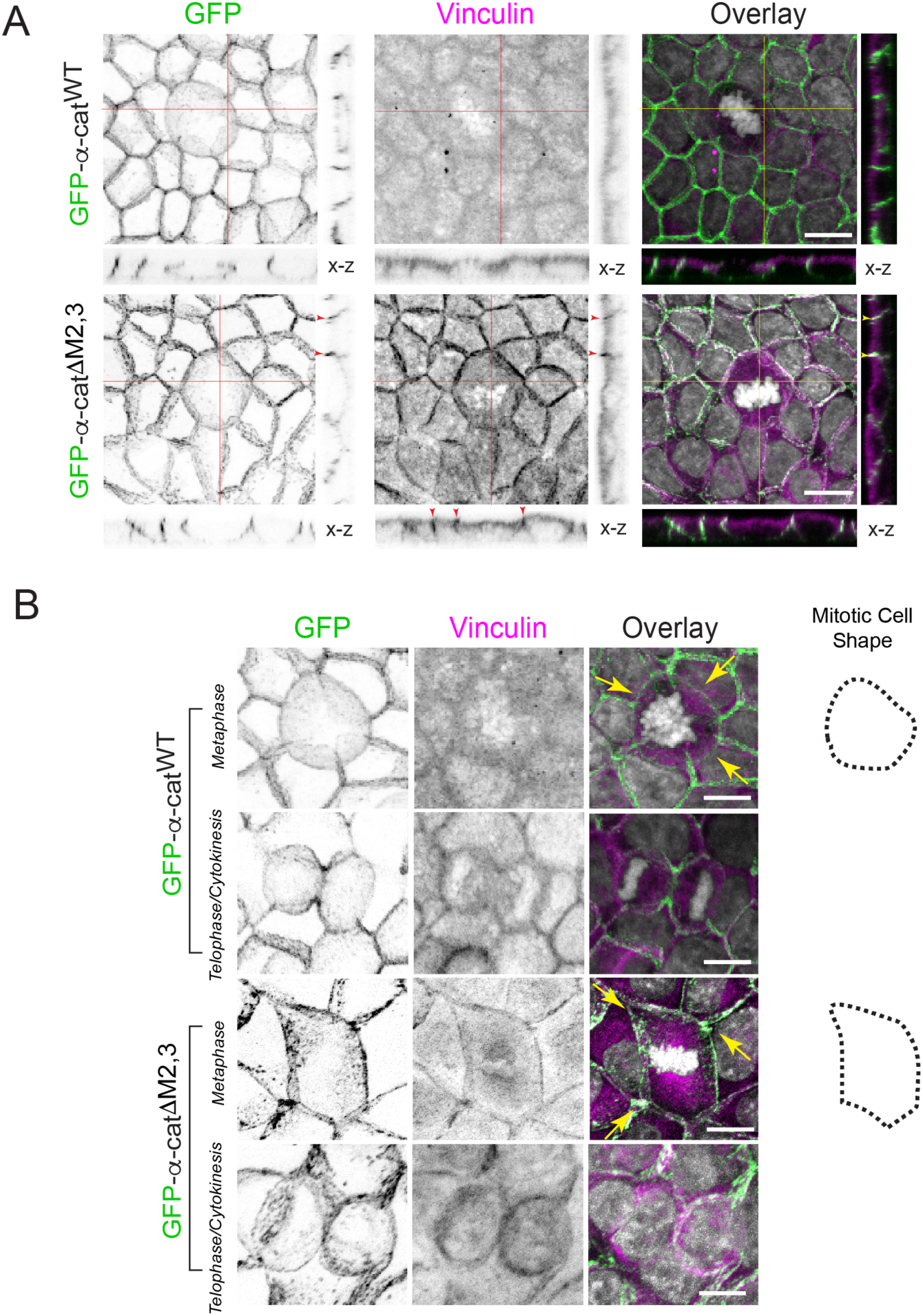
Persistent recruitment of vinculin to apical junctions via α-cat ΔM2-3 mutant impacts early mitotic rounding but not cytokinesis. **(A)** Confocal, maximum z-projection images (inverted grayscale) and x-z slices showing that α-cat-1′M2,3 constitutively and symmetrically recruits Vinculin to cell-cell junctions (red arrowheads), GFP-α-cat (green), LZTS2 (magenta). Scale bar, 5μm. **(B)** Confocal, maximum z-projection images (inverted grayscale) show that while α-cat-1′M2,3 limits mitotic rounding relative to α-cat-WT-expressing cells (yellow arrows; mitotic cell shape trace), binucleation rates are similar (see Fig. 10).

**Fig. S7:**
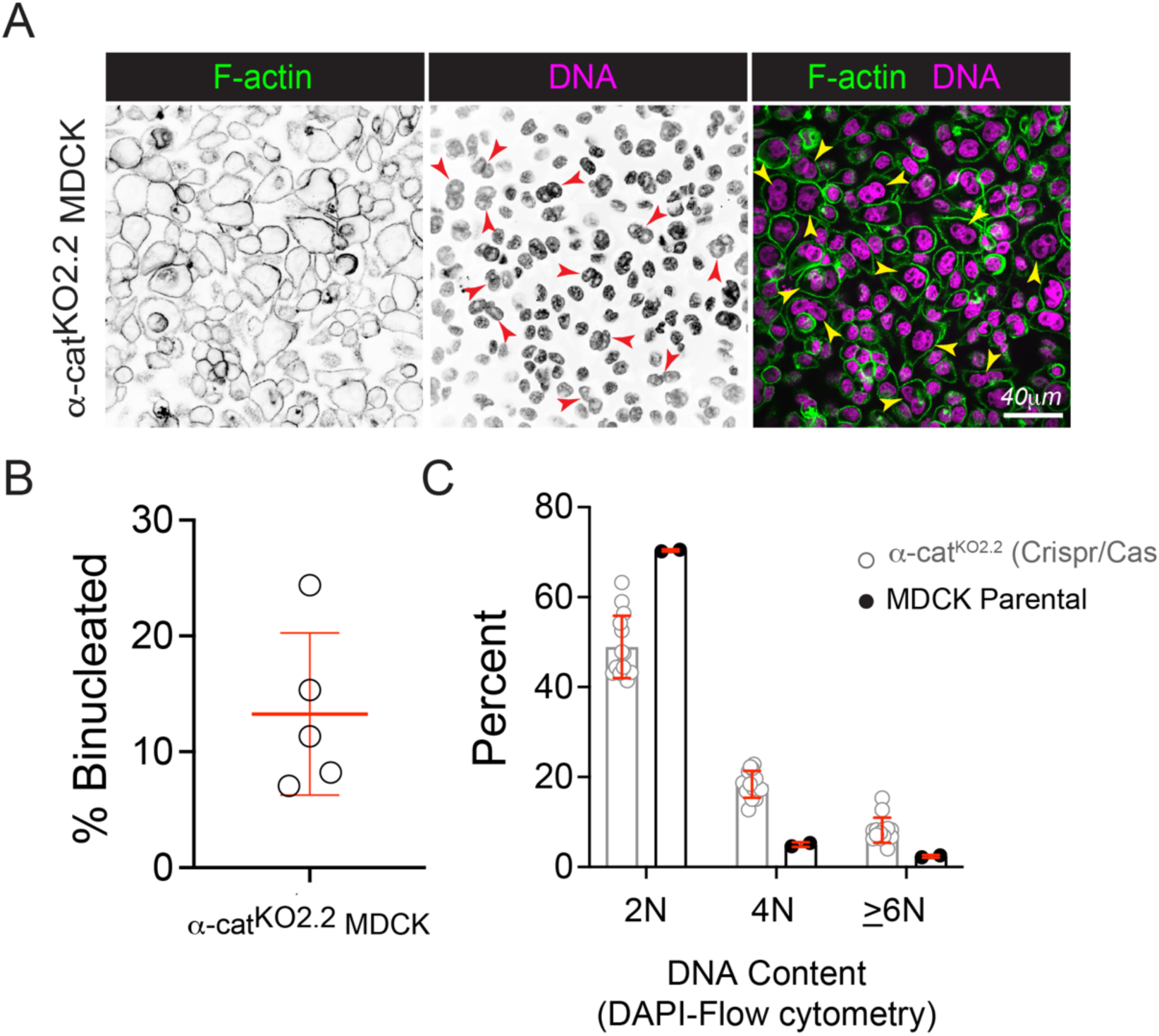
α-cat CRISPR knock-out MDCK cells show robust binucleation. **(A)** Confocal *en face* image showing binucleation is a prominent feature of α-cat CRISPR knock-out MDCK cells. Cellular junctions (Phalloidin, green), nuclei (Hoechst, magenta), and binucleated cells are indicated by red and yellow arrowheads. Scale bars, 40μm. (**B**) Quantification of α-cat^KO2.2^ MDCK binucleation rate. (**C**) DNA content of α-cat^KO2.2^ MDCK or MDCK parental cells measured by DAPI-staining and flow cytometry. Similar binucleation rates seen for other α-cat KO MDCK clones (not shown).

